# Global variability of the human IgG glycome

**DOI:** 10.1101/535237

**Authors:** Jerko Štambuk, Natali Nakić, Frano Vučković, Maja Pučić-Baković, Genadij Razdorov, Irena Trbojević-Akmačić, Mislav Novokmet, Toma Keser, Marija Vilaj, Tamara Pavić, Ivan Gudelj, Mirna Šimurina, Manshu Song, Hao Wang, Marijana Peričić Salihović, Harry Campbell, Igor Rudan, Ivana Kolčić, Leigh Anne Eller, Paul McKeigue, Merlin L. Robb, Jonas Halfvarson, Metin Kurtoglu, Vito Annese, Tatjana Škarić-Jurić, Mariam Molokhia, Ozren Polašek, Caroline Hayward, Hannah Kibuuka, Kujtim Thaqi, Dragan Primorac, Christian Gieger, Sorachai Nitayaphan, Tim Spector, Youxin Wang, Therese Tillin, Nish Chaturvedi, James F. Wilson, Moses Schanfield, Maxim Filipenko, Wei Wang, Gordan Lauc

**Affiliations:** Genos Glycoscience Research Laboratory, Zagreb, Croatia.; Department of Neuroscience, Scuola Internazionale Superiore di Studi Avanzati (SISSA), Trieste, Italy; Faculty of Pharmacy and Biochemistry, University of Zagreb, Zagreb, Croatia.; Beijing Key Laboratory of Clinical Epidemiology, School of Public Health, Capital Medical University, Beijing, China.; Institute for Anthropological Research, Zagreb, Croatia.; Centre for Global Health Research, Usher Institute of Population Health Sciences and Informatics, The University of Edinburgh, Edinburgh, United Kingdom.; School of Medicine, University of Split, Split, Croatia.; Walter Reed Army Institute of Research, Silver Spring, Maryland, USA.; Henry M. Jackson Foundation for the Advancement of Military Medicine, Bethesda, Maryland, USA.; Department of Gastroenterology, Faculty of Medicine and Health, Örebro University, Örebro, Sweden.; Department of Oncology, Koç University School of Medicine, Istanbul, Turkey.; Careggi University Hospital, Florence, Italy.; School of Population Health and Environmental Sciences, King’s College London, United Kingdom.; MRC Human Genetics Unit, MRC Institute for Genetics and Molecular Medicine, University of Edinburgh, Edinburgh, United Kingdom.; Makerere University Walter Reed Project, Kampala, Uganda.; Institute of Clinical Biochemistry, Priština, Kosovo.; Helmholtz Zentrum München - German Research Center for Environmental Health, Neuherberg, Germany.; Armed Forces Research Institute of Medical Sciences, Bangkok, Thailand.; Department of Twin Research & Genetic Epidemiology, King’s College London, London, United Kingdom.; Institute of Cardiovascular Science, Faculty of Population Health Sciences, London, United Kingdom.; Department of Forensic Sciences, George Washington University, Washington, DC, USA.; Institute of Chemical Biology and Fundamental Medicine, Novosibirsk, Russia.; School of Medical and Health Sciences, Edith Cowan University, Perth, Australia.

## Abstract

Immunoglobulin G (IgG) is the most abundant serum antibody and is a key determinant of the humoral immune response. Its structural characteristics and effector functions are modulated through the attachment of various sugar moieties called glycans. IgG N-glycome patterns change with the age of individual and in different diseases. Variability of IgG glycosylation within a population is well studied and is affected by a combination of genetic and environmental factors. However, global inter-population differences in IgG glycosylation have never been properly addressed. Here we present population-specific N-glycosylation patterns of whole IgG, analysed in 5 different populations totalling 10,482 IgG glycomes, and of IgG’s fragment crystallisable region (Fc), analysed in 2,530 samples from 27 populations sampled across the world. We observed that country of residence associates with many N-glycan features and is a strong predictor of monogalactosylation variability. IgG galactosylation also strongly correlated with the development level of a country, defined by United Nations health and socioeconomic development indicators. We found that subjects from developing countries had low IgG galactosylation levels, characteristic for inflammation and ageing. Our results suggest that citizens of developing countries may be exposed to country-specific environmental factors that can cause low-grade chronic inflammation and the apparent increase in biological age.

## Introduction

Immunoglobulin G is the most abundant glycoprotein and antibody class in human plasma^1^. It mediates interactions between antigens and the immune system^2^. There are four IgG subclasses present in plasma: IgG1, IgG2, IgG3 and IgG4^3^. Each subclass has distinctive functions, such as pronounced affinity for certain types of antigens, formation of immune complexes, complement activation, interactions with effector cells, half-life and placental transport^2^.

Glycosylation is co- and post-translational modification which is orchestrated by a complex biosynthetic pathway^4^. IgG contains a conserved N-glycosylation site on Asn297 residue within its fragment crystallisable (Fc) region on each of the two heavy chains^5^. Glycans attached to IgG are a complex biantennary type, with core structure consisting of four *N*-acetylglucosamines and three mannoses. Different glycan moieties such as bisecting GlcNAc, galactose, sialic acid and fucose can be attached to this core^6^. IgG shows a high degree of glycosylation diversity. Each of four IgG subclasses displays a distinctive glycan profile^7^. Also, each of the heavy chains of the same molecule can carry different glycans, creating a large repertoire of possible glycan patterns^8^. Finally, in 15-20% of cases, an additional N-glycosylation site appears within a variable region of the antibody, as a result of sequence variation in the variable region^9^.

The majority of IgG functions are achieved through interactions with receptors on immune cells and complement proteins. Fc glycans affect immunoglobulin conformation, which, in turn, defines binding affinity for Fc gamma receptors (FcγRs) on effector cells and complement, leading to alternations in effector functions^1,10^. IgG galactosylation has an extensive effect on its inflammatory potential^11^. Namely, agalactosylated IgG increases inflammation through activation of complement system^12,13^. Moreover, galactosylation was found necessary for C1q complement component binding and activation of complement-dependent cytotoxicity (CDC)^14^. It was also required for increased binding of IgG to activating Fc gamma receptors and therefore activation of antibody-dependent cellular cytotoxicity (ADCC)^15^. On the other hand, presence of galactose is necessary for activation of anti-inflammatory cascade through interactions with FcγRIIB and inhibition of the inflammatory activity of C5a complement component^16,17^. Presence of fucose attached to the first N-acetylglucosamine, i.e. core fucose, decreases ADCC activity, while the presence of bisecting GlcNAc increases binding to activating Fcγ receptors^18^. Terminal sialic acid increases anti-inflammatory roles of IgG by decreasing ADCC. Sialylated IgG molecules are also recognized by lectin receptors and complement component C1q, but proposed mechanisms underlying anti-inflammatory functions initiated through these interactions are still subject of debates^19^.

There is a prominent inter-individual variability of total IgG N-glycome^20^. Average IgG glycome heritability is approximately 50%, while the remaining variability can be mostly attributed to environmental factors^20,21,22^. The composition of IgG N-glycome gradually changes over the lifetime. It is strongly influenced by age, sex hormones and lifestyle^23^. Prominent changes in the IgG glycome were found in a number of diseases. In different autoimmune and alloimmune disorders, cancers and infectious diseases, IgG glycosylation changes reflect the increase inflammation, which is accompanying these conditions^11^. The impact of IgG glycosylation on its ability to modulate inflammation has been extensively studied as a potential biomarker for disease prognosis and therapy response, as well as for monoclonal antibody development^24,25^.

Ageing is a process of damage aggregation in an organism, leading to the disruption of health. It is influenced by both genetic factors and environment/lifestyle. In a healthy individual, a gender-specific gradual change in IgG glycosylation can be observed with an increase of chronological age. Namely, digalactosylated structures decrease with age, while agalactosylation and bisecting GlcNAc increase in older individuals^23^. On the other hand, changes in sialylation and core fucosylation displayed inconsistent trends in different studies^23^. Proposed model to describe and explain age-related changes in glycosylation is “the inflammageing model”, which implicates that inflammation causes changes in IgG glycosylation which, in turn, accelerate the process of ageing^26^. Consequently, changing under influence of both genes and environment, IgG glycans predict biological age and represent a measure of organisms health^11^.

Despite the fact that structural and functional aspects of IgG glycosylation are intensively studies and associated with predisposition and course of different diseases, little is known about the regulation of IgG glycosylation or mechanisms that lead to extensive changes in glycome composition after environmental challenge^11,27,28^. Therefore, the aim of this study was to estimate and compare various IgG N-glycosylation patterns in populations across the world as a result of their different genetic backgrounds and specific environmental influences.

## Results

### Total IgG glycans change with chronological age

In the initial analysis, samples originating from 10,482 healthy individuals and 5 different populations were analysed. Fluorescently labelled N-glycans released from IgG were separated into 24 chromatographic peaks (Supplementary Table 1; Supplementary Figure 1). Additionally, derived glycan traits (galactosylation, core fucosylation, sialylation and presence of bisecting GlcNAc) were calculated, based on the initial 24 glycan measures. Derived glycan traits represent a portion of structurally similar glycan species in the total IgG glycome (Supplementary Table 2). In general, galactosylation showed the highest variability of all IgG glycan traits, which is in line with the previous studies (Supplementary Figure 2).

It is known that the chronological age of subject affects IgG glycosylation^23^. Age-related changes were observed in the levels of IgG glycan traits in all studied populations (Figure 1a). Agalactosylated species and glycans containing bisecting *N*-acetylglucosamine (GlcNAc) increased with the chronological age of the participant. The opposite trend was observed for the levels of digalactosylated and sialylated glycans, which were decreasing with chronological age. On the other hand, core fucosylation and monogalactosylation levels did not change consistently with age. Also, age-related changes in glycosylation displayed sex-specific patterns, where female participants displayed more rapid changes.

**Figure 1:**
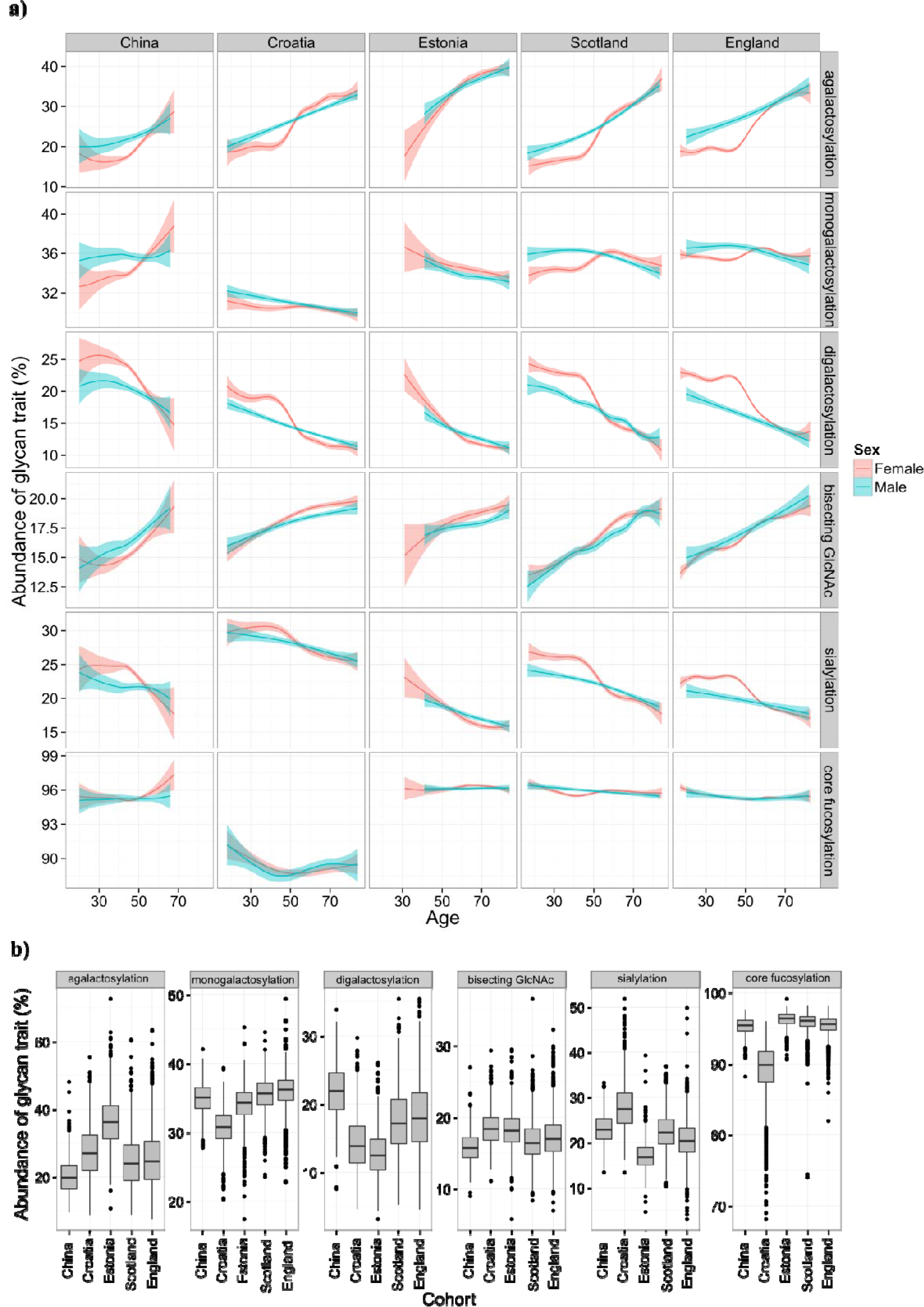
IgG glycan levels in five different populations. a, Relationship between age and derived glycan trait. Plots describe associations between each of five glycan traits and chronological age of participant. Blue and red curves represent fitted linear regression models. The shaded region is the 95% confidence interval on the fitted values. b, Differences in IgG glycosylation of participants from five populations. Each box represents interquartile range (25^th^ to 75^th^ percentiles). Lines inside the boxes represent the median values, while lines outside the boxes represent the 10^th^ and 90^th^ percentiles. Circles indicate outliers.

### Age and country of residence can predict total IgG glycosylation

Although total IgG N-glycans showed similar age-related changes within each of the studied cohorts, every population displayed particular glycan patterns (Figure 1b). Again, the most pronounced differences between populations were observed in the levels of agalactosylated glycans, which increased with a median age of the analysed population. This glycan trait had the lowest median value in young Chinese cohort (20%), while the highest was observed in Estonian cohort (36%), which contained the oldest population. Besides agalactosylation, pronounced differences between populations were also observed in the levels of digalactosylated and sialylated glycans (Supplementary Table 3). Relations of age, country of residence and sex with the total IgG glycans were evaluated to further investigate changes in IgG glycan traits in different populations. Chronological age appeared to be a good predictor of digalactosylation and agalactosylation variability (explaining 30% and 31% respectively), while it was able to describe 20% of bisecting GlcNAc variability. Contrary to age, participant’s country of residence appeared to be the strongest predictor of core fucose levels (*P*<6×10^−350^, n=5), explaining 57% of the variability in this glycan trait. It was also a good predictor of monogalactosylation and sialylation variability. On the other hand, sex was able to explain less than 1% of the variability of any glycan trait (Supplementary Table 4).

### Fc IgG glycan patterns in 14 countries

To validate observed diversity and unambiguously determine IgG N-glycosylation patterns in different populations, while eliminating potential batch effects, we compared glycan features derived from IgG subclass-specific Fc glycopeptides from 2,530 individuals (Supplementary Table 5, Supplementary Figure 3). This part of the study included 27 populations collected in 14 different countries (Supplementary Table 6). Subclass-specific glycopeptides were separated and accurate masses were measured for each glycoform. Calculated IgG Fc N-glycan derived traits displayed considerable variability between analysed populations. The Fc N-glycome composition is known to differ from the total IgG N-glycome, as a result of Fab N-glycome contribution to the total IgG glycome^29^. Again, the most prominent variation appeared to be in level of IgG1 galactosylation (Figure 2), although expected decrease of this glycan trait with the age of population was not observed. On the contrary, some populations appeared to have lower galactosylation than expected for the given chronological age. Population from Papua New Guinea, as the youngest one, surprisingly had the highest median level of agalactosylation (45%), while the subjects from England exhibited the lowest levels of this glycan trait (28%) on IgG1 subclass. The opposite effect was observed for monogalactosylation levels - the subjects from Papua New Guinea had the lowest median value of this glycan trait, while the highest levels were observed in the participants from England. In a similar manner, participants from countries such as Germany and Italy had higher monogalactosylation levels (comparable to subjects from England) than the ones from countries such as Uganda, which were more similar to the subjects from Papua New Guinea (Supplementary Table 7).

**Figure 2:**
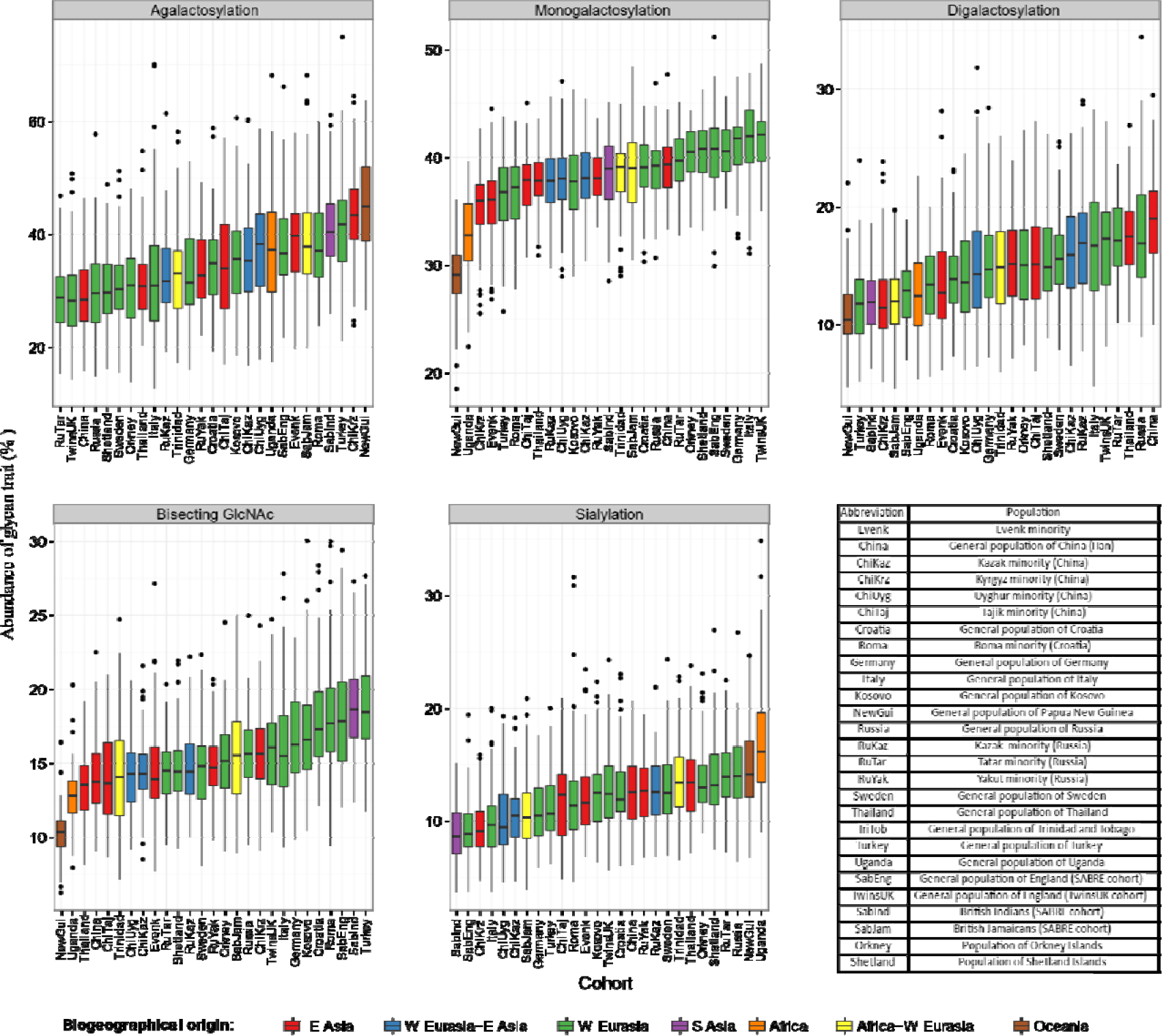
Levels of derived IgG1 glycan traits in 27 different populations. Each box represents interquartile range (25^th^ to 75^th^ percentiles) with median values drawn as middle line. Whiskers outside the boxes represent the 10^th^ and 90^th^ percentiles, while dots indicate outliers.

In the case of IgG2 and IgG4 subclasses, galactosylation levels displayed similar variation to IgG1 subclass, although observed glycosylation patterns appeared to be subclass-specific, especially in case of IgG4, which is the least abundant subclass in the human plasma (Supplementary Tables 8 and 9).

### Age and county of residence can predict IgG Fc glycosylation

To determine the source of variation in IgG Fc glycan profiles, we analysed relationship of glycan traits with sex, chronological age and country of residence. Here as well, chronological age was a predictor of agalactosylation and digalactosylation variability (Table 1). Interestingly, age was the best predictor of IgG2 digalactosylation. It was able to explain 28% of IgG2 agalactosylation variability compared to 22% in the case of IgG1.

**Table 1:** Proportion of glycan feature variability in 14 countries explained by linear mixed model with age and sex defined as fixed effects and country of residence as a random effect. Displayed values represent percentage (%) glycan trait variability explained by country, age and sex.

| IgG subclass | Glycan feature | Percent of glycan trait variability explained by country of residence (%) | Percent of glycan trait variability explained by age (%) | Percent of glycan trait variability explained by sex (%) | Country of origin *P* value |
| --- | --- | --- | --- | --- | --- |
| IgG1 | Agalactosylation | 21.4 | 18.3 | 0.9 | $1.0 \times 10^{-111}$ |
| | Monogalactosylation | 38.0 | 2.6 | 0.1 | $7.7 \times 10^{-193}$ |
| | Digalactosylation | 18.6 | 21.7 | 1.1 | $3.5 \times 10^{-98}$ |
| | Sialylation | 10.8 | 13.2 | 0.6 | $1.3 \times 10^{-54}$ |
| | Bisecting GlcNAc | 18.6 | 17.5 | 0.0 | $9.3 \times 10^{-110}$ |
| IgG2 | Agalactosylation | 12.8 | 23.9 | 0.9 | $7.2 \times 10^{-62}$ |
| | Monogalactosylation | 20.8 | 7.6 | 0.1 | $3.3 \times 10^{-94}$ |
| | Digalactosylation | 10.4 | 27.5 | 1.1 | $2.7 \times 10^{-54}$ |
| | Sialylation | 6.5 | 18.2 | 0.5 | $1.2 \times 10^{-27}$ |
| | Bisecting GlcNAc | 12.4 | 11.9 | 0.1 | $3.7 \times 10^{-64}$ |
| IgG4 | Agalactosylation | 20.5 | 13.7 | 0.4 | $1.3 \times 10^{-93}$ |
| | Monogalactosylation | 20.8 | 2.6 | 0.1 | $1.3 \times 10^{-78}$ |
| | Digalactosylation | 18.2 | 14.1 | 0.8 | $1.3 \times 10^{-86}$ |
| | Sialylation | 15.4 | 9.7 | 0.5 | $6.4 \times 10^{-75}$ |
| | Bisecting GlcNAc | 7.6 | 15.7 | 0.6 | $2.0 \times 10^{-37}$ |

Country of residence was the best predictor of IgG Fc monogalactosylation variability (Table 1). Namely, 38% of IgG1 Fc monogalactosylation variability was explained by the subject’s country of residence. Similar patterns were observed for IgG2 and IgG4 Fc glycosylation. The same as in the case of total IgG glycans, sex was able to explain up to 1% of the IgG Fc glycan variability. Therefore, chronological age and country of residence are good predictors of IgG Fc glycosylation.

### IgG Fc galactosylation correlates with the development level of a country

Development indicators measure the quality of specific life aspects. In order to resolve whether the observed associations between country of residence and studied IgG Fc glycan traits can be attributed to the development level of the country, we analysed relations between 45 development indicators and 5 derived glycan traits of each analysed IgG subclass (Supplementary Tables 10 - 12). The analysis resulted with 44 statistically significant correlations of IgG Fc monogalactosylation, digalactosylation and agalactosylation with 23 different development indicators. Majority of development indicators displayed significant positive correlations with IgG1 monogalactosylation (Supplementary Table 13). As for the subclasses IgG2 and IgG4, only monogalactosylation appeared to be significantly correlated with the studied indicators. On the other hand, we did not observe any significant correlations between any of the development indicators and sialylation or the incidence of bisecting GlcNAc on any of the IgG subclasses.

United Nation’s Human development index (HDI) is a summary measure of the development level of a certain country. It represents three dimensions of life: economy, education and health quality. Among the studied populations, Western European nations (Germany, England, Scotland, Sweden) have the highest development level expressed through HDI, while Papua New Guinea and Uganda have the lowest HDI scores. We found a positive correlation between HDI and IgG1 Fc monogalactosylation (Figure 3a), while it negatively correlated with IgG1 agalactosylation. These observations replicated on IgG2 subclass, where HDI positively correlated with monogalactosylation levels. Therefore, participants from developing countries appear to have lower IgG galactosylation when compared to their counterparts from developed countries.

**Figure 3:**
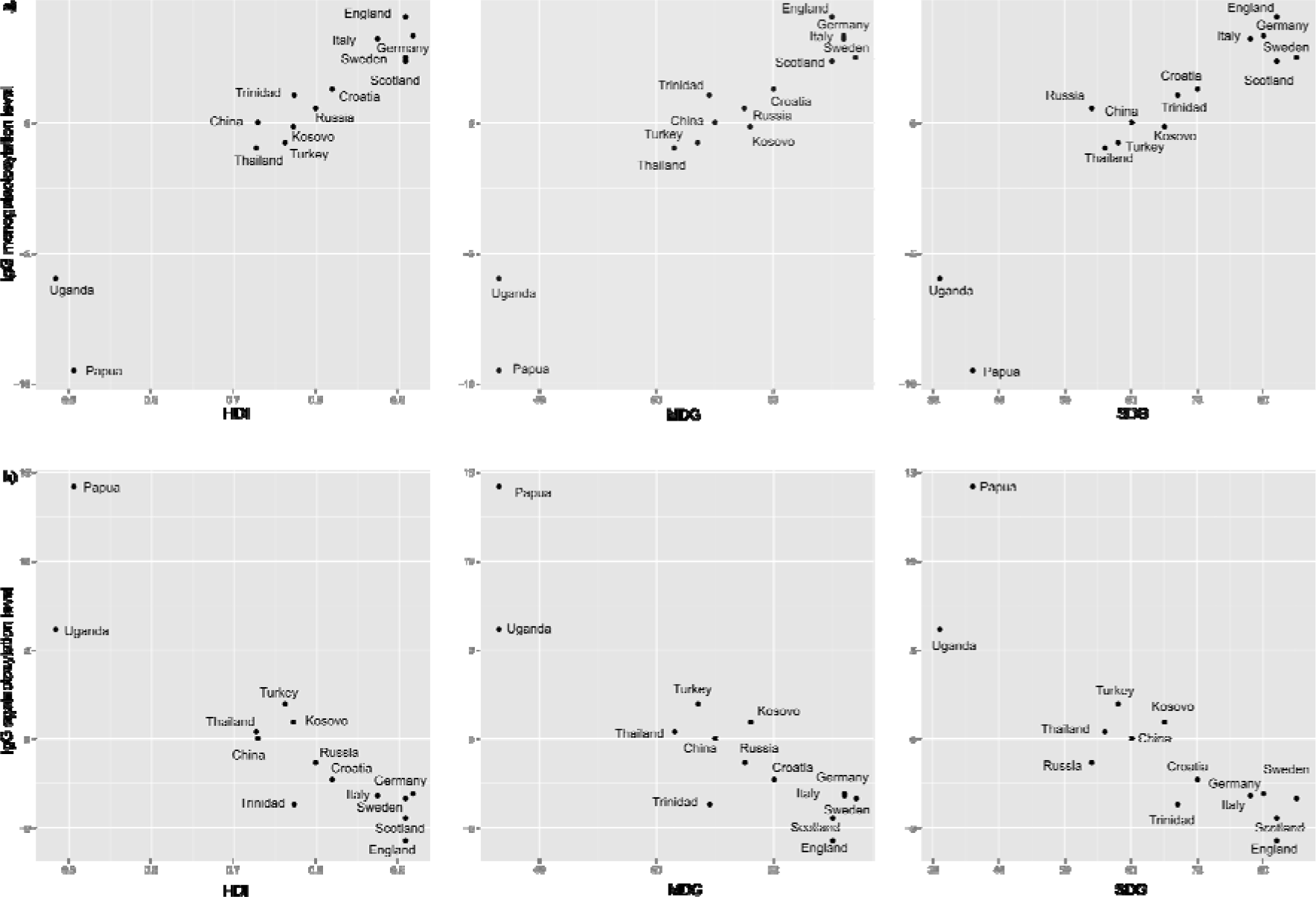
Relationship between IgG monogalactosylation level (a) and IgG agalactosylation level (b) with development indices in a specific country of residence. HDI = Human Development Index; SDI = health-related Sustainable Development Goals Index; MDG = health-related Millennium Development Goals Index.

### IgG Fc galactosylation as a marker of population’s health status

To determine the impact of the quality of health on IgG glycans, correlations between the two were calculated. Population’s health quality was expressed through overall health indices and specific health indicators. Countries with lower development level, in general, had also lower health-related indicators (Supplementary Table 12). Majority of health-related indicators appeared to be correlated with IgG monogalactosylation (Supplementary Table 13). Millennium development goals (MDG) index, which describes health-related indicators in MDG system, positively correlated with IgG1 and IgG2 monogalactosylation (r=0.97, *P*=7.44×10^−6^ and r=0.86, *P*=4.59×10^−2^ respectively) and IgG1 agalactosylation (r=−0.90, *P*=8.16×10^−3^; Figure 3a). In a similar fashion, positive correlation with IgG1 monogalactosylation displayed also the sustainable development goals (SDG) index, non-MDG index (health-related SDG indicators not included in MDG) and Health index, which like MDG index display overall health quality of a specific country (Supplementary Table 13, Figure 3a). SDG index was negatively correlated with IgG1 agalactosylation (Figure 3b).

Besides health-related indices, specific health-related indicators also correlated with IgG Fc galactosylation. Among all studied specific indicators, the decline in stunted growth prevalence demonstrated the strongest positive correlation with IgG1 monogalactosylation (r=0.97, *P*=1.16×10^−5^; *n*=14). Of the other studied indicators, universal health coverage and the decrease in occupational risk burden displayed substantial correlations with IgG Fc galactosylation. Life expectancy is also one of the most important indicators, used to describe life quality. Both female and male life expectancies were correlated with IgG Fc monogalactosylation. Exposure to various antigens was presented through indicators such as hygiene, water, WasH mortality and sanitation, which also displayed correlations with IgG Fc galactosylation. Of the infectious diseases, only hepatitis B showed a significant correlation with IgG Fc monogalactosylation (Supplementary Table 13).

Moreover, digalactosylation of IgG1 demonstrated five positive correlations with health-related development indicators, where skilled birth attendance and again, stunted growth, had the strongest associations with this glycan trait (Supplementary Table 13).

Although IgG Fc glycans showed the highest correlation coefficients with the health-related indicators, significant correlations between IgG1 Fc galactosylation features and the socioeconomic indicators such as education and economic development have also been determined. Education index was significantly correlated with both Fc monogalactosylation (r=0.94, *P*=0.0005, *n*=14) and agalactosylation (r=−0.90, *P*=0.010, *n*=14), while Gross Domestic Product (GDP) was significantly correlated only with monogalactosylation (r=0.89, *P*=0.01, *n*=14; Supplementary Table 13).

## Discussion

IgG N-glycosylation varies between individuals within the same population as well as between different populations^20,30^. In this study, we compared glycan profiles of the whole IgG molecule in 10 482 subjects originating from 5 different populations and Fc glycan profiles of 2 530 subjects from 27 cohorts and 25 ethnicities. Furthermore, this study yielded valuable data on IgG glycan levels in healthy participants from 14 countries and 25 ethnicities. The observed changes in glycome composition of the analysed populations suggest country-specificity of IgG glycan profiles.

Besides genetics, environment plays a crucial role in IgG glycosylation^28,22^. Pathogens, stress and nutrition are possible players orchestrating non-genomic component of the inter-populational variation in IgG glycosylation patterns^21,31^. Development level of a country reflects human well-being in a specific community and thereby environmental impact on an individual. We found that the development level of a country of residence was positively correlated with IgG galactosylation level. Namely, IgG galactosylation was decreased in people from countries with lower development level, while people from highly developed countries had also the highest levels of IgG galactosylation. These associations were observed on all IgG subclasses, although the largest number of development indicators correlated with glycans originating from IgG1 subclass, probably due to the highest concentration of this subclass in plasma.

Besides overall development level of a country, different health-related indicators displayed associations with IgG galactosylation level, indicating the impact of health quality on IgG glycosylation. IgG glycosylation modulates antibody’s pro- and anti-inflammatory actions, where a decline in galactosylation level, similar to the one observed in underdeveloped countries, was found in several inflammatory and autoimmune diseases, such as inflammatory bowel disease (IBD), rheumatoid arthritis and systemic lupus erythematosus^24,28,32^. Therefore, we speculated that populations with lower galactosylation have higher inflammatory potential of IgG and a higher low-grade systematic inflammation. Our findings are supported by a recent study on 773 children, which compared IgG glycosylation in subjects from Gabon, Ghana, Ecuador, the Netherlands and Germany. The increase in agalactosylated species was observed in individuals from Gabon, Ghana and Ecuador, compared to participants from Netherlands and Germany. These changes were correlated with the history of parasitic infections and generalised to immune activation^30^.

In our study, the prevalence of stunted growth displayed a strong correlation with glycosylation. This may be partially explained by the fact that stunted growth is caused by environmental enteropathy, which is a chronic intestinal inflammation caused by malnutrition, continuous bacterial exposure, repeated enteric infections and small intestinal bacterial overgrowth^33^. Since many inflammatory conditions are associated with decreased IgG galactosylation levels, environmental enteropathy could cause changes in IgG galactosylation by inducing chronic subclinical inflammation in the gastrointestinal tract^34^. Hence, our results imply that country-specific differences in the quality of health influence IgG inflammatory potential and cause specific glycosylation patterns in different countries.

Furthermore, we also observed significant correlations between the traits describing IgG Fc galactosylation and the socioeconomic indicators. Economy and education quality are reflected through country’s development level and quality of the health system^35^. For that reason, our findings are not surprising and further emphasize environmental influence on the health of the certain population.

IgG glycosylation is known to change with chronological age of individual^23^. Within five populations where the whole IgG glycan profiles were measured, we observed similar decrease in galactosylation levels with the age of participant. This age-related decline in IgG galactosylation is believed to be one of the causes of higher inflammation in older individuals^23,36^. Although exact mechanisms are still unclear, there are several proposed pathways which could explain underlying age-related changes in IgG glycosylation. Possible mechanisms include various expression and/or activity of glycosylation-related enzymes, selection of B-cell clones with specific IgG glycan patterns and B-cell independent glycosylation. Furthermore, a decrease in IgG galactosylation was observed in the premature ageing syndromes^36^. Through modulation of inflammation, IgG galactosylation, or, more precisely, agalactosylation is proposed to contribute to biological ageing in a process of inflammaging^37^. Since the decrease in IgG galactosylation is a hallmark of increased biological age, the proinflammatory IgG Fc glycosylation profile in individuals from developing countries may imply accelerated biological ageing in these populations, resulting in a shorter expected lifespan.

In summary, we revealed that at a community level, immunoglobulin G glycosylation patterns vary between different countries. We also correlated changes in galactosylation with participant’s chronological age and development level of a country. Constant environmental pressure on the immune system in developing countries maintains IgG constantly in under-galactosylated, proinflammatory state. As a consequence of this permanent low-degree IgG Fc galactosylation, individuals from developing countries display premature populational ageing and appear to be biologically “older” than residents of more developed countries.

## Materials and methods

### Study participants

Total IgG glycome analysis was based on 10,482 human participants from China^38,28^, Croatia^39^, Estonia and two cohorts from the United Kingdom (population of Scotland from Orkney Islands^20^ and England from the TwinsUK cohort^40^) (Supplementary Table 1). Subclass specific analysis of IgG Fc glycosylation included 2,530 healthy individuals. Volunteers originated from 14 different countries and 25 different ethnic groups (Supplementary Table 6). For Kazak and English cohorts, we had two populations obtained from different medical centres. Samples were randomized across 96-well plates (31 in total), with 5 technical replicates of a standard sample and 1 blank, serving as a negative control. Development level of a country was assessed using three development indices: health-related Sustainable Development Goal index (SDG)^41^, health-related Millennium Development Goal Index (MDG)^41^ and United Nation Human Development Index (HDI)^42^, while specific aspects of human life were assessed using other development indicators (Supplementary Table 13). Blocking was performed by equally distributing subjects of the same sex and age from all the cohorts across used plates. Plasma samples used as standards were obtained from Croatian National Institute of Transfusion Medicine. Study was performed in compliance with the Helsinki declaration and all participants gave written informed consent. Ethical approval was obtained by relevant ethics committees.

### Immunoglobulin G isolation

Protein G affinity chromatography was used to isolate immunoglobulin G from human blood plasma as described previously^20^. In short, maximum volume of 100 μL of human peripheral blood plasma or serum were diluted with 1X phosphate buffer saline (PBS) and loaded onto protein G monolithic plate (BIA Separations, Ajovščina, Slovenia). Samples were washed three times with 1X PBS and IgG was eluted using 0.1M formic acid (Merck, Darmstadt, Germany) followed by immediate neutralisation with 1M ammonium bicarbonate (Acros Organics, Pittsburgh, PA).

### Immunoglobulin G trypsin digestion and purification

IgG glycopeptides were obtained and purified as described before^43^. Approximately 15 μg of isolated IgG was treated with 0.1 μg of sequencing grade trypsin (Promega, Fitchburg, WI) and incubated overnight at 37 °C. Reaction was stopped by dilution with 0.1% trifluoroacetic acid (TFA; Sigma-Aldrich, St. Louis, MI). Glycopeptides were purified using a solid phase extraction on Chromabond C-18 sorbent (Macherey-Nagel, Düren, Germany). Samples were loaded onto beads in 0.1% TFA, and washed three times using the same solvent. Glycopeptides were eluted from the phase with 20% LC-MS grade acetonitrile (ACN; Honeywell, Morris Plains, NJ). Eluted glycopeptides were vacuum-dried and reconstituted in 20 μL of ultrapure water prior to LC-MS analysis. All glycan analyses were performed at Genos laboratory.

### Release and labelling of the total IgG N-glycans

Glycan release and labelling of Croatian samples was performed as previously described^23^. Briefly, IgG was incorporated into sodium dodecyl sulphate polyacrylamide gel and glycans were released from protein using an overnight incubation with PNGase F (ProZyme, Hayward, CA). Released glycans were labelled with 2-aminobenzamide (2-AB; Sigma-Aldrich) and purified on Whatman 3 mm chromatography paper. For cohorts from Scotland, England, China and Estonia, glycans were released as previously described^23,31^. Briefly, IgG was denatured using 1.33% (w/v) sodium dodecyl sulphate (Invitrogen, Carlsbad, CA) and samples were incubated at 65 °C for 10 minutes. Subsequently, 4% (v/v) Igepal CA-630 (Sigma–Aldrich) and 1.25 mU of PNGase F (ProZyme) were added to each sample and incubated overnight at 37 °C. For glycan labelling, 48 mg/mL of 2-AB in dimethyl sulfoxide (Sigma–Aldrich) and glacial acetic acid (Merck) (v/v 85:15) was mixed with reducing agent (106.96 mg/mL of 2-picoline borane (Sigma–Aldrich) in dimethyl sulfoxide). Labelling mixture was added to samples, followed by 2-hour incubation at 65 °C.

After incubation, Estonian and Chinese samples were brought to 96% ACN (J.T. Baker, Phillipsburg, NJ) and applied to each well of a 0.2 μm GHP filter plate (Pall Corporation, Ann Arbor, MI). Samples were subsequently washed five times using acetonitrile/water (96:4, v/v). Glycans were eluted with water and stored at −20 °C until usage. Samples from England and Scotland were purified using a solid-phase extraction on 200 μL of 0.1 g/L microcrystalline cellulose suspension (Merck) in a 0.45 μm GHP filter plate (Pall Corporation). Deglycosylation reaction was diluted four times with ACN loaded to cellulose. Samples were washed three times with 80% ACN and eluted with ultrapure water.

### HILIC-UPLC analysis of fluorescently labelled N-glycans

Fluorescently labelled N-glycans were separated by hydrophilic interaction liquid chromatography (HILIC) on a Waters Acquity UPLC H-class instrument (Waters, Milford, MA) equipped with FLR fluorescence detector set to 330 nm for excitation and 420 nm for emission wavelength. Separation was achieved on a Waters ethylene bridged hybrid (BEH) Glycan chromatography column, 100 × 2.1 mm i.d., 1.7 μm BEH particles with 100 mM ammonium formate (pH 4.4) as a solvent A and ACN as a solvent B. Separation method used linear gradient from 75% to 62% solvent B (v/v) at a flow rate of 0.4 mL/min in a 25-minute analytical run. Column temperature was maintained at 60 °C. Obtained chromatograms were manually separated into 24 peaks from which, using the total area normalisation, relative abundances of 24 directly measured glycan traits were obtained.

### LC-MS analysis of IgG Fc glycopeptides

Trypsin-digested, subclass-specific glycopeptides were separated and measured on nanoAcquity chromatographic system (Waters, Milford, MA) coupled to Compact mass spectrometer (Bruker, Bremen, Germany), equipped with Apollo II source as described previously with minor changes^44^. Samples (9 μL) were loaded onto PepMap 100 C8 trap column (5 mm × 300 μm i.d.; Thermo Fisher Scientific, Waltham, MA) at a flow rate of 40 μL/min of solvent A (0.1% TFA) and washed of salts and impurities for one minute. Subclass-specific glycopeptides were separated on C18 analytical column (150 mm × 100 μm i.d., 100 Å; Advanced Materials Technology, Wilmington, DE) in a gradient from 18% to 25% of solvent B (80% ACN) in solvent A. Column temperature was set to 30 °C and flow rate was 1 μl/min. NanoAcquity was coupled to mass spectrometer via capillary electrophoresis sprayer interface (Agilent, Santa Clara, CA), which allows mixing of analytical flow with sheath liquid (50% isopropanol, 20% propionic acid; Honeywell, Morris Plains, NJ).

Mass spectrometer was operated in a positive ion mode, with capillary voltage set to 4500 V, nebulizer pressure set to 0.4 bar and drying gas set to 4 l/min at 180 °C. Spectra were recorded in a *m/z* range of 600 - 1800. Collision energy was 4 eV.

Obtained raw data was converted to centroid mzXML files using ProteoWizard version 3.0.1. software. Samples were internally calibrated using defined list of IgG glycopeptides with highest signal-to-noise ratios and required isotopic patterns. After calibration, signals matching IgG Fc glycopeptides were extracted from data using 10 *m/z* extraction window. First four isotopic peaks of doubly and triply charged signals, belonging to the same glycopeptide species, were summed together, resulting in 20 glycopeptides per IgG subclass. Predominant allotype variant of IgG3 tryptic peptide carrying N-glycans in Caucasian population has the same amino acid sequence as IgG2. On the other hand, in Asian and African populations predominant variant of the same peptide has the same amino acid composition as IgG4 making the separation of IgG3 from other subclasses impossible using given separation methods^45^. Therefore, IgG glycopeptides were separated into three chromatographic peaks labeled IgG1, IgG2 and IgG4. Signals of interest were normalised to the total area of each IgG subclass.

### Statistical analysis

Data analysis was performed using program R, version 3.0.1. with a ggplot2 package for creation of visualisations. Derived glycan traits representing levels of galactosylation (agalactosylation, monogalactosylation and digalactosylation), sialylation, core fucosylation and incidence of bisecting GlcNAc were calculated from obtained data as described before (Supplementary Tables 2 and 5)^43^. In short, traits were calculated as portion of glycans containing common structural features in a total IgG glycome, or in a specific subclass of IgG Fc glycopeptides. Core fucosylation was excluded from IgG Fc specific glycopeptide analysis due to low data quality of non-fucosylated species. To remove experimental and batch variation, batch correction was performed using a R ComBat package followed by log-transformation of glycan or glycopeptide data. Linear mixed model was used to analyse correlations between glycan traits and the subject’s country of residence. In the model, sex and age were fixed effects, while the country of residence was used as a random effect. Likelihood ratio test was used to determine significance of country of origin variability in glycan trait variability. Pearson’s correlation coefficient was used to express relationships between country-specific development indicator and level of glycan trait in participants from the same country. *P* values were adjusted for multiple testing using Bonferroni correction.

### Data availability

The data that support the findings of this study are available from the corresponding author upon reasonable request.

## Acknowledgements

This study was supported by European Structural and Investment Funds IRI grant (#KK.01.2.1.01.0003) and Croatian National Centre of Research Excellence in Personalized Healthcare grant (#KK.01.1.1.01.0010). Collection of Chinese cohorts was supported by National Natural Science Foundation of China (81573215, 81773527, 81673247 and 81370083) and Australian-China Collaborative Grant (NH&MRC-APP1112767–NSFC 81561128020).

## Author contributions

G.L. designed the study. J.Š., N.N., G.R. and M.N. carried out LC-MS analysis. F.V. performed statistical analysis. M.P.B., I.T.A., T.K., T.P., M.Š., I.G., and M.V. performed UPLC analysis. M.S., H.W., Y.W., W.W., M.P.S., T.Š.J., H.C., C.H., J.F.W., I.R., O.P., I.K., S.N., L.A.E., H.K., M.L.R., M.M., P.M., J.H., M.K., V.A., K.T., C.G., T.S., T.T., N.C., M.S. and M.F. recruited participants and provided plasma samples. J.Š. drafted the manuscript. All authors edited and approved the final version.

## Conflict of interest

GL is founder and CEO of Genos – a private research organization that specializes in high-throughput glycomic analysis and has several patents in this field. J.Š., F.V., M.P.B., G.R., I.T.A., I.G., M.V., and M.N. are employees of Genos.

Correspondence and requests for materials should be addressed to G.L.

