## Supplementary Figures for "Global variability of the human IgG glycome"

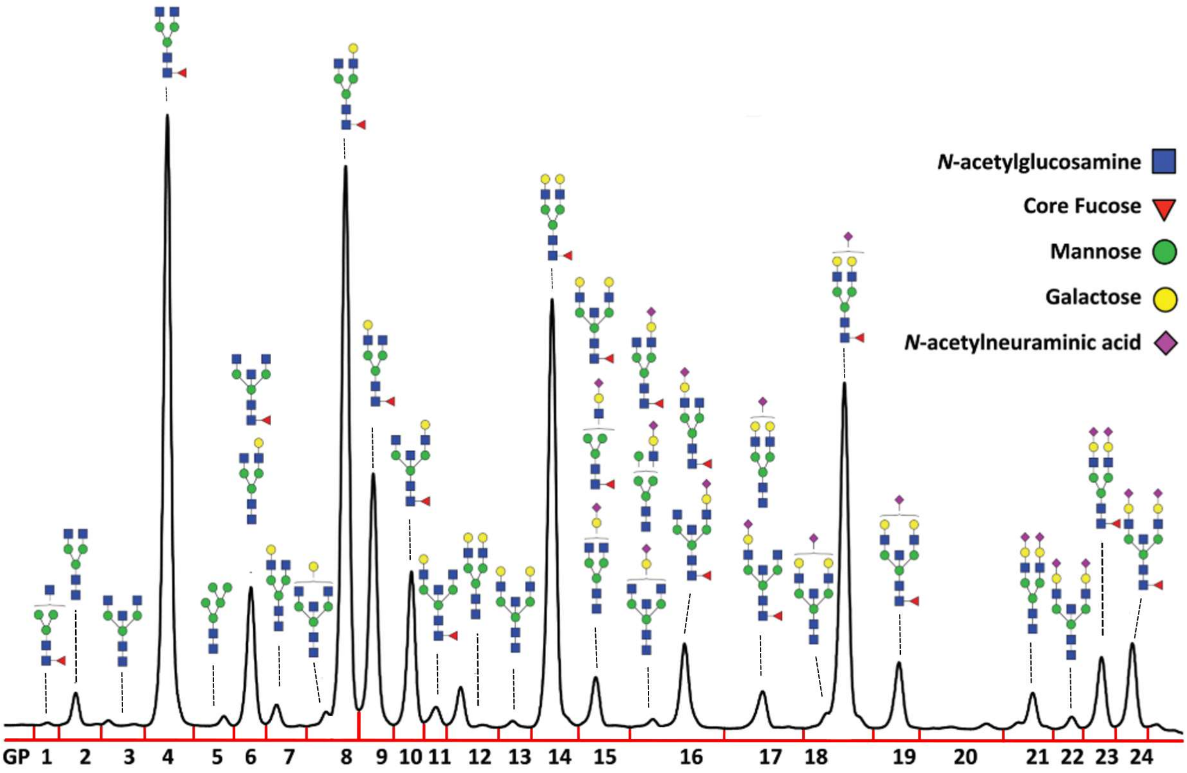


Supplementary Figure: 1 HILIC-UPLC chromatogram of total IgG released glycans labelled with 2-AB. Dominant structures are indicated above each of 24 glycan peaks (GP 1 - 24).

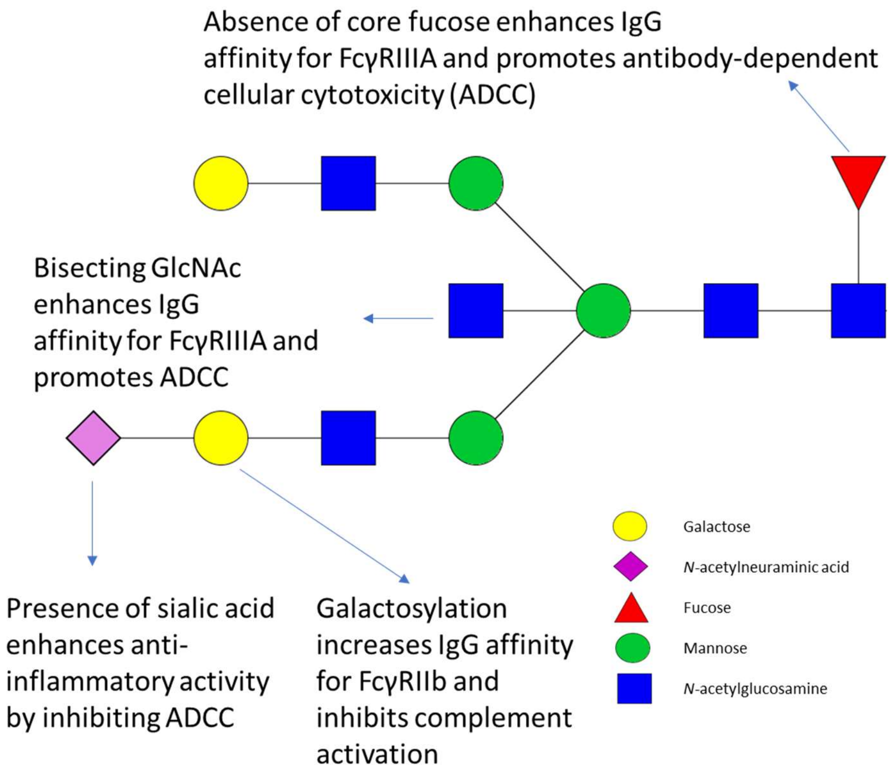


Supplementary Figure 2: Effects of alternative glycan moieties on IgG Fc region. Glycans influence structure of IgG Fc region and manipulate Fc effector functions[44].

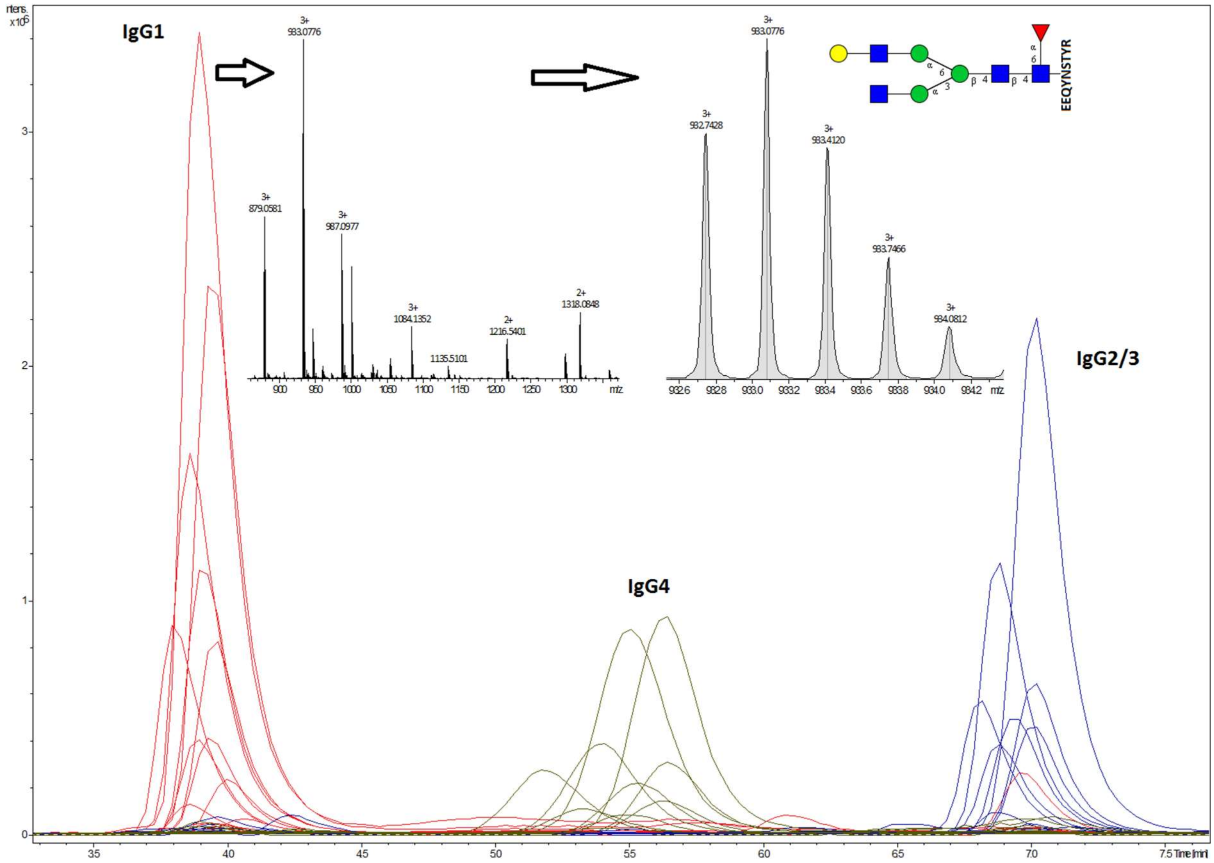


Supplementary Figure 3: Extracted ion chromatograms of IgG Fc glycopeptides. IgG1 subclass is shown in red, IgG4 in green and IgG2 in blue. Mass spectra containing masses that correspond to IgG1 glycopeptides and isotopic distribution of the most abundant glycopeptide (G1F) are shown.
