## Supplementary Tables for "Global variability of the human IgG glycome"

Supplementary Table 1: Demographic characteristics of five populations used for HILIC-UPLC analysis of total IgG glycosylation. In table are given: abbreviation of analysed population, country of participant's residence, age parameters (minimum, maximum, median, mean, 1$^{st}$ and 3$^{rd}$ quartile) and sex parameters (number of female and male participants).

| Population | Abbreviation | Country of residence | Min age | Q1 | Median | Mean | Q3 | Max age | F | M |
|---|---|---|---|---|---|---|---|---|---|---|
| General population of China (Han) | China | China | 20 | 43 | 48 | 47 | 51 | 68 | 423 | 201 |
| General population of Croatia | Croatia | Croatia | 18 | 46 | 57 | 56 | 68 | 98 | 1116 | 689 |
| General population of Estonia | Estonia | Estonia | 31 | 60 | 69 | 67 | 75 | 88 | 516 | 593 |
| Population of Orkney Islands | Scotland | Scotland | 17 | 41 | 54 | 53 | 65 | 100 | 1236 | 794 |
| General population of England (TwinsUK cohort) | England | England | 17 | 42 | 54 | 52 | 62 | 83 | 4282 | 342 |

Supplementary Table 2: Description of directly measured and derived IgG glycan traits measured by HILIC-UPLC.

| Glycan peak | The most abundant glycan structure[1] | Desctiption of glycan trait | Trait calculation[2] |
|---|---|---|---|
| GP1 | 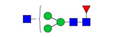 | *The percentage of FA1 glycan in total IgG glycans* | *GP1 / TOTAL GLYCANS* 100* |
| GP2 | 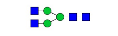 | *The percentage of A2 glycan in total IgG glycans* | *GP2 / TOTAL GLYCANS* 100* |
| GP3 | 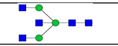 | *The percentage of A2B glycan in total IgG glycans* | *GP3 / TOTAL GLYCANS* 100* |
| GP4 | 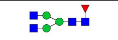 | *The percentage of FA2 glycan in total IgG glycans* | *GP4 / TOTAL GLYCANS* 100* |
| GP5 | 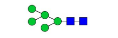 | *The percentage of M5 glycan in total IgG glycans* | *GP5 / TOTAL GLYCANS* 100* |
| GP6 | 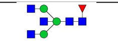 | *The percentage of FA2B glycan in total IgG glycans* | *GP6 / TOTAL GLYCANS* 100* |
| GP7 | 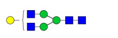 | *The percentage of A2G1 glycan in total IgG glycans* | *GP7 / TOTAL GLYCANS* 100* |
| GP8 | 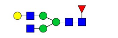 | *The percentage of FA2[6]G1 glycan in total IgG glycans* | *GP8 / TOTAL GLYCANS* 100* |
| GP9 | 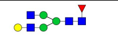 | *The percentage of FA2[3]G1 glycan in total IgG glycans* | *GP9 / TOTAL GLYCANS* 100* |
| GP10 | 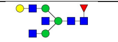 | *The percentage of FA2[6]BG1 glycan in total IgG glycans* | *GP10 / TOTAL GLYCANS* 100* |
| GP11 | 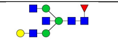 | *The percentage of FA2[3]BG1 glycan in total IgG glycans* | *GP11 / TOTAL GLYCANS* 100* |
| GP12 | 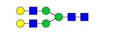 | *The percentage of A2G2 glycan in total IgG glycans* | *GP12 / TOTAL GLYCANS* 100* |
| GP13 | 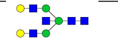 | *The percentage of A2BG2 glycan in total IgG glycans* | *GP13 / TOTAL GLYCANS* 100* |
| GP14 | 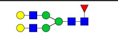 | *The percentage of FA2G2 glycan in total IgG glycans* | *GP14 / TOTAL GLYCANS* 100* |
| GP15 | 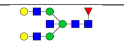 | *The percentage of FA2BG2 glycan in total IgG glycans* | *GP15 / TOTAL GLYCANS* 100* |
| GP16 | 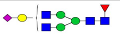 | *The percentage of FA2G1S1 glycan in total IgG glycans* | *GP16 / TOTAL GLYCANS* 100* |
| GP17 | 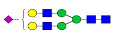 | *The percentage of A2G2S1 glycan in total IgG glycans* | *GP17/ TOTAL GLYCANS* 100* |
| GP18 | 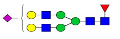 | *The percentage of FA2G2S1 glycan in total IgG glycans* | *GP18 / TOTAL GLYCANS* 100* |
| GP19 | 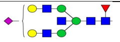 | *The percentage of FA2BG2S1 glycan in total IgG glycans* | *GP19 / TOTAL GLYCANS* 100* |
| GP20 | | *Structure not determined* | *GP20 / TOTAL GLYCANS* 100* |
| GP21 | 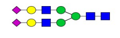 | *The percentage of A2G2S2 glycan in total IgG glycans* | *GP21 / TOTAL GLYCANS* 100* |
| GP22 | 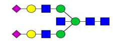 | *The percentage of A2BG2S2 glycan in total IgG glycans* | *GP22 / TOTAL GLYCANS* 100* |
| GP23 | 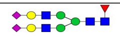 | *The percentage of FA2G2S2 glycan in total IgG glycans* | *GP23 / TOTAL GLYCANS* 100* |
| GP24 | 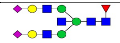 | *The percentage of FA2BG2S2 glycan in total IgG glycans* | *GP24 / TOTAL GLYCANS* 100* |

| Derived glycan trait | Derived trait description | Derived trait calculation |
|---|---|---|
| Core fucosylation | Fraction of structures containing core fucose in total IgG glycans | GP1+GP4+GP6+GP8+GP9+GP10+GP11+GP14+GP15+GP16+GP18+GP19+GP23+GP24 |
| Bisecting GlcNAc | Fraction of structures containing bisecting GlcNAc in total IgG glycans | GP3+GP6+GP10+GP11+GP13+GP15+GP19+GP22+GP24 |
| Agalactosylation | Fraction of agalactosylated structures in total IgG glycans | GP1+GP2+GP3+GP4+GP5+GP6 |
| Monogalactosylation | Fraction of structures containing one galactose in total IgG glycans | GP7+GP8+GP9+GP10+GP11+GP16 |
| Digalactosylation | Fraction of structures containing two galactoses in total IgG glycans | GP12+GP13+GP14+GP15+GP17+GP18+GP19+GP20+GP21+GP22+GP23+GP24 |
| Sialylation | Fraction of structures containing sialic acid in total IgG glycans | GP16+GP17+GP18+GP19+GP20+GP21+GP22+GP23+GP24 |

[1]Glycan structures are drawn in GlycoWorkbench version 2. blue square = N-acetylglucosamine, red triangle = fucose, green circle = mannose, yellow circle = galactose, purple diamond = N-acetylneuraminic acid.

[2]Total glyacans = sum of all 24 glycan peaks

Supplementary Table 3: Derived glycan traits in five populations used for HILIC-UPLC analysis of IgG glycosylation. Mean values of agalactosylation, monogalactosylation, digalactosylation, bisecting GlcNAc, sialylation and core fucosylation are given with minimal and maximal values in brackets.

| Population | Agalactosylation %, (min - max) | Monogalactosylation %, (min - max) | Digalactosylation %, (min - max) | Bisecting GlcNAc %, (min - max) | Sialylation %, (min - max) | Core fucosylation %, (min - max) |
|---|---|---|---|---|---|---|
| China | 20.0 (16.5 - 23.5) | 35.0 (33.4 - 36.4) | 21.9 (19.1 - 24.6) | 15.6 (14.2 - 17.2) | 22.9 (20.6 - 25.4) | 95.4 (94.6 - 96.1) |
| Croatia | 27.1 (21.9 - 32.5) | 30.8 (29.1 - 32.4) | 13.8 (11.3 - 16.7) | 18.4 (16.8 - 20.0) | 27.3 (24.3 - 30.9) | 89.9 (87.5 - 91.7) |
| Estonia | 36.2 (31.3 - 41.1) | 34.3 (32.6 - 35.8) | 12.4 (10.3 - 14.8) | 18.2 (16.6 - 19.9) | 16.7 (15.0 - 18.8) | 96.4 (95.7 - 96.9) |
| Scotland | 24.0 (18.9 - 29.7) | 35.6 (34.0 - 37.1) | 17.1 (14.1 - 20.7) | 16.5 (14.9 - 18.5) | 22.2 (19.6 - 25.1) | 96.1 (95.3 - 96.6) |
| England | 24.6 (19.2 - 30.5) | 36.2 (34.7 - 37.6) | 17.9 (14.5 - 21.6) | 17.0 (15.2 - 18.9) | 20.4 (17.9 - 23.2) | 95.6 (94.7 - 96.3) |

Supplementary Table 4: Linear mixed model in five populations with age and sex defined as fixed effects and country of residence as a random effect. Displayed values represent percentage (%) glycan trait variability explained by age, sex and country of residence, with country of residence likelihood-ratio test *P* value.

| Derived glycan trait | Percent of glycan trait variability explained by country of residence (%) | Percent of glycan trait variability explained by age (%) | Percent of glycan trait variability explained by sex (%) | Country of residence *P* value |
|---|---|---|---|---|
| Agalactosylation | 14.4 | 30.9 | 0.8 | $6.6 \times 10^{-194}$ |
| Monogalactosylation | 39.9 | 0.0 | 0.2 | $< 6 \times 10^{-350}$ |
| Digalactosylation | 19.5 | 30.2 | 0.7 | $1.1 \times 10^{-301}$ |
| Bisecting GlcNAc | 6.3 | 20.1 | 0.1 | $6.7 \times 10^{-104}$ |
| Sialylation | 42.5 | 11.9 | 0.2 | $< 6 \times 10^{-350}$ |
| Core fucosylation | 57.5 | 0.1 | 0.0 | $< 6 \times 10^{-350}$ |

Supplementary Table 5: Description of directly measured and derived subclass-specific Fc IgG glycan traits measured by LC-MS with mass list.

| Glycan trait[1] | Glycan scheme[2] | IgG1 glycopeptide m/z[3] | | IgG2&3 glycopeptide m/z[4] | | IgG4glycopeptide m/z[5] | | Glycan trait description | Glycan trait calculation[6] |
|---|---|---|---|---|---|---|---|---|---|
| | | $[M+2H]^{2+}$ | $[M+3H]^{3+}$ | $[M+2H]^{2+}$ | $[M+3H]^{3+}$ | $[M+2H]^{2+}$ | $[M+3H]^{3+}$ | | |
| G0F | 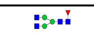 | 1317.527 | 878.687 | 1301.532 | 868.024 | 1309.529 | 873.356 | Fraction of FA2 glycan in total subclass Fc glycans | *G0F/total subclass Fc glycans*100* |
| G1F | 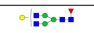 | 1398.553 | 932.705 | 1382.558 | 922.042 | 1390.556 | 927.373 | Fraction of FA2G1 glycan in total subclass Fc glycans | *G1F/total subclass Fc glycans*100* |
| G2F | 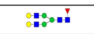 | 1479.58 | 986.722 | 1463.585 | 976.059 | 1471.582 | 981.391 | Fraction of FA2G2 glycan in total subclass Fc glycans | *G2F/total subclass Fc glycans*100* |
| G0FN | 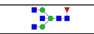 | 1419.067 | 946.38 | 1403.072 | 935.717 | 1411.069 | 941.049 | Fraction of FA2B glycan in total subclass Fc glycans | *G0FN/total subclass Fc glycans*100* |
| G1FN | 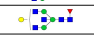 | 1500.093 | 1000.398 | 1484.098 | 989.735 | 1492.096 | 995.066 | Fraction of FA2BG1 glycan in total subclass Fc glycans | *G1FN/total subclass Fc glycans*100* |
| G2FN | 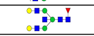 | 1581.119 | 1054.416 | 1565.125 | 1043.752 | 1573.122 | 1049.084 | Fraction of FA2BG2 glycan in total subclass Fc glycans | *G2FN/total subclass Fc glycans*100* |
| G1FS1 | 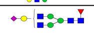 | 1544.101 | 1029.737 | 1528.106 | 1019.073 | 1536.104 | 1024.405 | Fraction of FA2G1S1 glycan in total subclass Fc glycans | *G1FS1/total subclass Fc glycans*100* |
| G2FS1 | 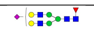 | 1625.127 | 1083.754 | 1609.133 | 1073.091 | 1617.13 | 1078.423 | Fraction of FA2G2S1 glycan in total subclass Fc glycans | *G2FS1/total subclass Fc glycans*100* |
| G1FNS1 | 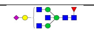 | 1645.641 | 1097.430 | 1629.646 | 1086.767 | 1637.643 | 1092.098 | Fraction of FA2BG1S1 glycan in total subclass Fc glycans | *G1FNS1/total subclass Fc glycans*100* |
| G2FNS1 | 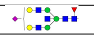 | 1726.667 | 1151.447 | 1710.672 | 1140.784 | 1718.67 | 1146.116 | Fraction of FA2BG2S1 glycan in total subclass Fc glycans | *G2FNS1/total subclass Fc glycans*100* |
| G0 | 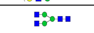 | 1244.498 | 830.001 | 1228.503 | 819.338 | 1236.501 | 824.67 | Fraction of A2 glycan in total subclass Fc glycans | *G0/total subclass Fc glycans*100* |
| G1 | 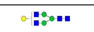 | 1325.524 | 884.019 | 1309.529 | 873.356 | 1317.527 | 878.687 | Fraction of A2G1 glycan in total subclass Fc glycans | *G1/total subclass Fc glycans*100* |
| G2 | 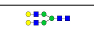 | 1406.551 | 938.036 | 1390.556 | 927.373 | 1398.553 | 932.705 | Fraction of A2G2 glycan in total subclass Fc glycans | *G2/total subclass Fc glycans*100* |
| G0N | 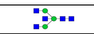 | 1346.038 | 897.694 | 1330.043 | 887.031 | 1338.04 | 892.363 | Fraction of A2B glycan in total subclass Fc glycans | *G0N/total subclass Fc glycans*100* |
| G1N | 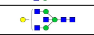 | 1427.064 | 951.712 | 1411.069 | 941.049 | 1419.067 | 946.38 | Fraction of A2BG1 glycan in total subclass Fc glycans | *G1N/total subclass Fc glycans*100* |
| G2N | 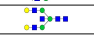 | 1508.090 | 1005.730 | 1492.096 | 995.066 | 1500.093 | 1000.398 | Fraction of A2BG2 glycan in total subclass Fc glycans | *G2N/total subclass Fc glycans*100* |
| G1S1 | 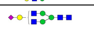 | 1471.072 | 981.051 | 1455.077 | 970.387 | 1463.075 | 975.719 | Fraction of A2G1S1 glycan in total subclass Fc glycans | *G1S1/total subclass Fc glycans*100* |
| G2S1 | 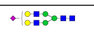 | 1552.098 | 1035.068 | 1536.104 | 1024.405 | 1544.101 | 1029.737 | Fraction of A2G2S1 glycan in total subclass Fc glycans | *G2S1/total subclass Fc glycans*100* |
| G1NS1 | 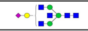 | 1572.612 | 1048.744 | 1556.617 | 1038.081 | 1564.614 | 1043.412 | Fraction of A2BG1S1 glycan in total subclass Fc glycans | *G1NS1/total subclass Fc glycans*100* |
| G2NS1 | 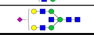 | 1653.638 | 1102.761 | 1637.643 | 1092.098 | 1645.641 | 1097.43 | Fraction of A2BG2S1 glycan in total subclass Fc glycans | *G2NS1/total subclass Fc glycans*100* |

| Derived glycan trait | Derived trait description | Derived trait calculation |
|---|---|---|
| Core fucosylation | Fraction of structures containing core fucose in subclass specific Fc glycans | G0F+G1F+G2F+G0FN+G1FN+G2FN+G1FS1+G2FS1+G1FNS1+G2FNS1 |
| Bisecting GlcNAc | Fraction of structures containing bisecting GlcNAc in subclass specific Fc glycans | G0FN+G1FN+G2FN+G1FNS1+G2FNS1+G0N+G1N+G2N+G1NS1+G2NS1 |
| Agalactosylation | Fraction of agalactosylated structures in subclass specific Fc glycans | G0+G0F+G0N+G0FN |
| Monogalactosylation | Fraction of structures containing one galactose in subclass specific Fc glycans | G1F+G1FN+G1FS1+G1FNS1+G1+G1N+G1S1+G1NS1 |
| Digalactosylation | Fraction of structures containing two galactoses in subclass specific Fc glycans | G2F+G2FN+G2FS1+G2FNS1+G2+G2N+G2S1+G2NS1 |
| Sialylation | Fraction of structures containing sialic acid in subclass specific Fc glycans | G1FS1+G2FS1+G1FNS1+G2FNS1+G1S1+G2S1+G1NS1+G2NS1 |

[1]Glycan composition: N (N-acetylglucosamine), F (fucose), G (galactose) and S (N-acetylneuraminic acid) followed by a number represents number and type of monosaccharides attached to A2 glycan.

[2]Glycan structures are drawn in GlycoWorkbench version 2. blue square = N-acetylglucosamine, red triangle = fucose, green circle = mannose, yellow circle = galactose, purple diamond = N-acetylneuraminic acid.

[3] IgG1 tryptic peptide sequence carrying glycan: $E_{293}EQYNSTYR_{301}$

[4] IgG4 tryptic peptide sequence carrying glycan: $E2_{93E}QFNSTFR_{301}$

[5] IgG2&3 tryptic peptide sequence carrying glycan: $E_{293}EQFNSTYR_{301}$

[6]total subclass Fc glycans = sum of all 20 glycopeptides in one IgG subclass

Supplementary Table 6: Demographic characteristics of 27 populations used for IgG Fc glycosylation analysis. In table are given: country of residence of participants, abbreviation of analysed cohort, age parameters (minimum, maximum, median, mean, 1st and 3rd quartile) and sex parameters (number of female and male participants).

| Population | Country of residence | Cohort abbreviation | Min. | Q1 | Median | Mean | Q3 | Max. | F | M |
|---|---|---|---|---|---|---|---|---|---|---|
| Evenk minority | China | Evenk | 10 | 22 | 29 | 33 | 41 | 85 | 36 | 74 |
| General population of China (Han) | China | China | 20 | 26 | 44 | 40 | 51 | 60 | 57 | 43 |
| Kazak minority | China | ChiKaz | 21 | 33 | 45 | 44 | 56 | 78 | 52 | 45 |
| Kyrgyz minority | China | ChiKrz | 22 | 55 | 65 | 62 | 70 | 92 | 56 | 41 |
| Uyghur minority | China | ChiUyg | 8 | 38 | 49 | 47 | 59 | 85 | 64 | 35 |
| Tajik minority | China | ChiTaj | 18 | 34 | 42 | 44 | 54 | 72 | 52 | 44 |
| General population of Croatia | Croatia | Croatia | 18 | 38 | 49 | 50 | 62 | 81 | 25 | 71 |
| Roma minority | Croatia | Roma | 18 | 28 | 37 | 38 | 42 | 79 | 54 | 58 |
| General population of Germany | Germany | Germany | 32 | 43 | 47 | 49 | 57 | 70 | 33 | 64 |
| General population of Italy | Italy | Italy | 24 | 34 | 46 | 47 | 58 | 86 | 48 | 39 |
| General population of Kosovo | Kosovo | Kosovo | 21 | 33 | 46 | 44 | 57 | 62 | 69 | 50 |
| General population of Papua New Guinea | Papua New Guinea | NewGui | 3 | 8 | 12 | 18 | 26 | 48 | 47 | 22 |
| General population of Russia | Russia | Russia | 9 | 17 | 20 | 23 | 26 | 43 | 74 | 26 |
| Kazak minority | Russia | RuKaz | 21 | 34 | 42 | 43 | 53 | 63 | 62 | 38 |
| Tatar minority | Russia | RuTar | 18 | 23 | 31 | 31 | 36 | 48 | 57 | 40 |
| Yakut minority | Russia | RuYak | 20 | 37 | 47 | 47 | 56 | 75 | 68 | 13 |
| General population of Sweden | Sweden | Sweden | 22 | 33 | 44 | 42 | 51 | 64 | 42 | 54 |
| Uninfected HIV cohort participants from Thailand | Thailand | Thailand | 18 | 28 | 32 | 33 | 36 | 49 | 56 | 66 |
| General population of Trinidad and Tobago | Trinidad and Tobago | TriTob | 22 | 40 | 53 | 50 | 59 | 77 | 82 | 14 |
| General population of Turkey | Turkey | Turkey | 16 | 48 | 59 | 57 | 67 | 79 | 39 | 61 |
| General population of Uganda | Uganda | Uganda | 17 | 29 | 33 | 33 | 37 | 47 | 61 | 50 |
| General population of England (SABRE cohort) | England | SabEng | 59 | 63 | 67 | 68 | 72 | 77 | 18 | 78 |
| General population of England (TwinsUK cohort) | England | TwinsUK | 21 | 33 | 46 | 46 | 57 | 70 | 48 | 47 |
| British Indians (SABRE cohort) | England | SabInd | 60 | 65 | 69 | 69 | 73 | 83 | 13 | 85 |
| British Jamaicans (SABRE cohort) | England | SabJam | 59 | 66 | 69 | 69 | 73 | 81 | 60 | 36 |
| Population of Orkney Islands | Scotland | Orkney | 21 | 33 | 46 | 45 | 56 | 70 | 49 | 50 |
| Population of Shetland Islands | Scotland | Shetland | 21 | 34 | 46 | 45 | 56 | 69 | 52 | 46 |

Supplementary Table 7: Derived glycan traits in 27 populations used for IgG1 Fc glycopeptide analysis. Median levels of agalactosylation, monogalactosylation, digalactosylation, bisecting GlcNAc, and sialylation with minimal and maximal values in brackets are given. Population abbreviations are defined in Figure 2 and Supplementary Table 6.

| Subclass | Population | Agalactosylation | Monogalactosylation | Digalactosylation | Bisecting GlcNAc | Sialylation |
|---|---|---|---|---|---|---|
| IgG1 | China | 28.5 (24.7 - 33.7) | 39.2 (37.1 - 40.9) | 19.0 (16.1 - 21.3) | 13.8 (12.3 - 15.7) | 12.6 (10.1 - 14.8) |
| IgG1 | ChiKaz | 35.3 (29.9 - 41.1) | 38.0 (36.1 - 40.4) | 15.8 (13.0 - 19.1) | 14.3 (13.3 - 15.6) | 10.5 (8.5 - 12.0) |
| IgG1 | ChiKrz | 43.3 (39.1 - 47.9) | 36.0 (33.7 - 37.4) | 11.4 (9.7 - 13.8) | 15.6 (13.7 - 17.3) | 9.2 (7.7 - 11.0) |
| IgG1 | ChiUyg | 37.8 (30.6 - 43.6) | 38.0 (36.1 - 39.8) | 14.3 (11.4 - 17.9) | 14.2 (12.3 - 15.7) | 9.4 (7.9 - 12.4) |
| IgG1 | Croatia | 34.9 (29.3 - 39.1) | 39.0 (37.1 - 41.1) | 13.8 (11.7 - 15.8) | 17.3 (15.4 - 19.8) | 11.9 (10.8 - 14.4) |
| IgG1 | Evenk | 39.4 (32.8 - 43.7) | 36.1 (33.7 - 37.8) | 12.7 (10.5 - 16.2) | 13.9 (12.6 - 16.1) | 11.6 (9.6 - 14.0) |
| IgG1 | Germany | 31.5 (27.6 - 39.2) | 41.8 (39.1 - 42.8) | 14.7 (12.3 - 17.6) | 16.3 (14.4 - 19.2) | 10.5 (8.6 - 12.9) |
| IgG1 | Italy | 31.0 (24.7 - 38.0) | 41.9 (39.2 - 44.3) | 16.7 (12.7 - 20.4) | 15.5 (13.4 - 18.2) | 9.7 (7.7 - 11.4) |
| IgG1 | Kosovo | 35.6 (29.5 - 40.6) | 37.7 (35.1 - 40.2) | 13.6 (11.0 - 17.1) | 16.6 (14.6 - 18.9) | 12.5 (9.9 - 14.1) |
| IgG1 | NewGui | 44.9 (38.9 - 51.9) | 29.0 (27.2 - 30.8) | 10.3 (8.8 - 12.5) | 10.3 (9.4 - 11.1) | 14.1 (11.9 - 17.0) |
| IgG1 | Orkney | 31.0 (25.2 - 35.8) | 40.5 (38.6 - 42.3) | 15.0 (12.1 - 18.0) | 15.1 (13.3 - 17.0) | 13.0 (11.6 - 14.9) |
| IgG1 | Roma | 36.9 (31.7 - 43.6) | 37.2 (34.1 - 39.1) | 13.4 (10.9 - 15.8) | 17.7 (15.8 - 20.1) | 11.3 (9.1 - 13.5) |
| IgG1 | RuKaz | 31.7 (27.9 - 37.4) | 37.8 (35.8 - 39.9) | 16.8 (13.4 - 19.5) | 14.4 (12.8 - 16.3) | 12.6 (10.6 - 14.9) |
| IgG1 | Russia | 29.4 (24.3 - 34.8) | 39.2 (37.0 - 40.6) | 16.9 (14.0 - 20.9) | 15.6 (14.0 - 17.2) | 13.9 (11.9 - 16.5) |
| IgG1 | RuTar | 28.8 (24.4 - 32.5) | 39.7 (37.8 - 41.7) | 17.2 (15.0 - 20.0) | 14.5 (12.9 - 15.8) | 13.9 (12.1 - 16.0) |
| IgG1 | RuYak | 32.7 (28.8 - 39.1) | 38.0 (36.4 - 40.0) | 15.1 (12.4 - 18.0) | 14.7 (13.5 - 16.1) | 12.7 (10.4 - 14.7) |
| IgG1 | SabEng | 36.6 (32.8 - 42.8) | 40.7 (37.9 - 42.6) | 12.9 (10.6 - 14.5) | 17.8 (15.1 - 20.5) | 8.9 (7.7 - 10.6) |
| IgG1 | SabInd | 40.4 (36.2 - 45.4) | 38.8 (36.0 - 41.0) | 11.9 (10.0 - 13.7) | 18.7 (16.7 - 20.7) | 8.6 (7.1 - 10.7) |
| IgG1 | SabJam | 37.8 (33.0 - 43.8) | 39.0 (35.7 - 41.3) | 12.0 (9.8 - 13.9) | 15.5 (12.9 - 17.8) | 10.3 (8.5 - 12.5) |
| IgG1 | Shetland | 29.7 (26.0 - 34.8) | 40.8 (38.6 - 42.4) | 14.9 (12.9 - 18.2) | 14.4 (13.2 - 16.1) | 13.1 (11.4 - 15.9) |
| IgG1 | Sweden | 30.3 (26.8 - 34.6) | 40.6 (38.5 - 42.5) | 15.6 (13.4 - 17.6) | 14.8 (12.6 - 16.2) | 12.5 (10.6 - 15.0) |
| IgG1 | Thailand | 31.1 (26.8 - 34.9) | 37.7 (36.3 - 39.5) | 17.4 (14.8 - 19.5) | 13.5 (11.8 - 14.8) | 13.4 (10.9 - 15.5) |
| IgG1 | TriTob | 33.1 (26.9 - 37.1) | 39.1 (36.7 - 40.4) | 14.8 (11.7 - 17.8) | 14.1 (11.5 - 16.5) | 13.4 (11.3 - 15.7) |
| IgG1 | Turkey | 41.8 (35.2 - 46.0) | 36.7 (34.0 - 39.0) | 11.7 (9.2 - 13.8) | 18.5 (16.7 - 20.9) | 10.6 (9.2 - 13.1) |
| IgG1 | Ugand | 37.3 (29.8 - 43.9) | 32.7 (30.2 - 35.7) | 12.4 (9.9 - 15.2) | 12.8 (11.6 - 13.8) | 16.2 (13.4 - 19.6) |
| IgG1 | TwinsUK | 28.3 (23.8 - 32.8) | 42.0 (39.6 - 43.3) | 17.3 (13.4 - 19.5) | 16.1 (13.5 - 17.7) | 12.4 (10.0 - 14.9) |
| IgG1 | ChiTaj | 34.0 (26.9 - 41.7) | 37.8 (35.5 - 39.3) | 15.1 (12.2 - 18.3) | 13.6 (11.6 - 16.4) | 12.4 (8.7 - 14.1) |

Supplementary Table 8: Derived glycan traits in 27 populations used for IgG2 Fc glycopeptide analysis. Median levels of agalactosylation, monogalactosylation, digalactosylation, bisecting GlcNAc, and sialylation with minimal and maximal values in brackets are given. Population abbreviations are defined in Figure 2 and Supplementary Table 6.

| Subclass | Population | Agalactosylation | Monogalactosylation | Digalactosylation | Bisecting GlcNAc | Sialylation |
|---|---|---|---|---|---|---|
| IgG2 | China | 37.3 (32.5 - 43.6) | 34.8 (32.9 - 36.5) | 15.9 (12.9 - 18.4) | 10.8 (9.5 - 12.1) | 11.4 (9.4 - 13.3) |
| IgG2 | ChiKaz | 40.6 (34.8 - 47.3) | 33.0 (30.9 - 34.6) | 14.6 (11.7 - 17.7) | 11.1 (9.8 - 12.2) | 11.7 (9.8 - 13.5) |
| IgG2 | ChiKrz | 51.4 (44.9 - 55.0) | 29.7 (27.9 - 32.1) | 10.4 (8.9 - 12.7) | 11.4 (10.5 - 13.1) | 9.1 (7.6 - 11.5) |
| IgG2 | ChiUyg | 44.6 (37.3 - 49.5) | 31.5 (29.7 - 33.6) | 12.4 (10.0 - 15.9) | 10.8 (9.5 - 11.9) | 10.9 (8.5 - 14.1) |
| IgG2 | Croatia | 43.7 (39.3 - 49.4) | 33.1 (30.5 - 34.8) | 11.7 (10.0 - 14.0) | 12.3 (10.5 - 14.6) | 11.2 (9.5 - 12.7) |
| IgG2 | Evenk | 43.8 (38.3 - 50.3) | 32.4 (30.0 - 34.1) | 12.7 (9.6 - 15.1) | 10.9 (9.8 - 12.5) | 9.9 (7.6 - 11.8) |
| IgG2 | Germany | 41.1 (35.3 - 47.3) | 34.6 (32.1 - 36.8) | 12.8 (10.6 - 15.6) | 12.2 (10.1 - 13.6) | 11.5 (9.2 - 13.0) |
| IgG2 | Italy | 37.8 (30.9 - 44.0) | 36.4 (32.7 - 38.5) | 14.9 (12.2 - 18.8) | 12.2 (10.7 - 13.5) | 11.2 (7.8 - 13.1) |
| IgG2 | Kosovo | 42.2 (36.0 - 49.7) | 31.9 (29.1 - 33.7) | 12.3 (9.8 - 15.2) | 12.2 (11.1 - 13.9) | 12.2 (10.1 - 14.6) |
| IgG2 | NewGui | 48.8 (43.0 - 52.2) | 28.4 (26.5 - 30.4) | 11.9 (9.8 - 13.8) | 9.5 (8.5 - 10.2) | 12.0 (9.5 - 14.7) |
| IgG2 | Orkney | 42.2 (35.2 - 48.7) | 33.2 (31.5 - 35.3) | 12.6 (9.8 - 15.5) | 11.2 (10.0 - 12.5) | 12.3 (10.0 - 14.0) |
| IgG2 | Roma | 42.6 (38.2 - 50.1) | 31.8 (29.0 - 33.9) | 12.4 (9.8 - 14.8) | 13.3 (11.3 - 14.7) | 12.3 (8.9 - 14.2) |
| IgG2 | RuKaz | 37.9 (34.4 - 44.8) | 33.2 (31.0 - 34.9) | 15.0 (12.0 - 17.4) | 11.1 (9.6 - 12.5) | 11.7 (10.1 - 14.0) |
| IgG2 | Russia | 36.9 (31.1 - 42.1) | 33.2 (31.2 - 35.2) | 15.6 (13.2 - 18.3) | 11.9 (10.7 - 13.0) | 13.8 (11.9 - 16.7) |
| IgG2 | RuTar | 36.2 (30.7 - 40.6) | 34.4 (32.3 - 36.0) | 16.0 (14.0 - 18.0) | 10.9 (10.0 - 12.2) | 13.8 (11.2 - 15.9) |
| IgG2 | RuYak | 39.1 (35.4 - 48.6) | 33.1 (31.2 - 35.7) | 14.5 (11.8 - 16.4) | 11.3 (10.4 - 12.8) | 11.0 (8.9 - 13.1) |
| IgG2 | SabEng | 46.1 (42.0 - 51.4) | 32.9 (30.4 - 35.7) | 10.5 (8.9 - 12.1) | 13.8 (12.1 - 16.0) | 9.9 (8.3 - 11.1) |
| IgG2 | SabInd | 48.7 (44.0 - 52.5) | 31.6 (29.1 - 33.3) | 10.2 (8.5 - 12.1) | 14.3 (13.3 - 16.2) | 9.5 (7.9 - 11.3) |
| IgG2 | SabJam | 45.6 (41.8 - 52.8) | 32.1 (30.0 - 34.5) | 9.9 (7.9 - 12.2) | 12.4 (10.9 - 14.3) | 10.2 (8.1 - 12.6) |
| IgG2 | Shetland | 41.1 (34.7 - 45.6) | 34.0 (32.4 - 36.2) | 12.8 (10.5 - 15.7) | 11.0 (9.9 - 12.2) | 11.8 (9.9 - 14.5) |
| IgG2 | Sweden | 40.1 (35.5 - 45.2) | 34.4 (33.0 - 36.1) | 13.3 (11.3 - 15.8) | 11.0 (9.6 - 12.3) | 12.0 (9.9 - 14.2) |
| IgG2 | Thailand | 34.0 (30.5 - 37.8) | 34.2 (32.7 - 35.3) | 17.0 (14.9 - 19.0) | 10.4 (9.1 - 12.0) | 14.4 (12.4 - 16.2) |
| IgG2 | TriTob | 42.4 (34.3 - 47.4) | 33.8 (31.6 - 35.9) | 12.5 (10.1 - 16.1) | 10.4 (8.5 - 12.0) | 11.3 (9.1 - 13.8) |
| IgG2 | Turkey | 47.9 (42.1 - 53.1) | 31.1 (28.7 - 33.1) | 10.3 (8.5 - 13.1) | 13.5 (12.0 - 15.2) | 10.4 (8.7 - 12.1) |
| IgG2 | Ugand | 41.2 (33.9 - 48.5) | 31.2 (28.8 - 32.9) | 12.8 (10.5 - 16.1) | 9.5 (8.7 - 11.3) | 14.0 (11.7 - 16.6) |
| IgG2 | TwinsUK | 36.3 (30.7 - 43.5) | 35.0 (32.3 - 36.8) | 14.5 (11.6 - 17.4) | 11.4 (10.1 - 13.3) | 13.6 (9.7 - 16.1) |
| IgG2 | ChiTaj | 41.3 (33.6 - 49.3) | 33.4 (30.1 - 34.8) | 14.3 (11.2 - 18.0) | 11.1 (9.7 - 12.8) | 11.1 (7.7 - 13.4) |

Supplementary Table 9: Derived glycan traits in 27 populations used for IgG4 Fc glycopeptide analysis. Median levels of agalactosylation, monogalactosylation, digalactosylation, bisecting GlcNAc, and sialylation with minimal and maximal values in brackets are given. Population abbreviations are defined in Figure 2 and Supplementary Table 6.

| Subclass | Population | Agalactosylation | Monogalactosylation | Digalactosylation | Bisecting GlcNAc | Sialylation |
|---|---|---|---|---|---|---|
| IgG4 | China | 33.5 (26.9 - 40.1) | 33.5 (30.9 - 37.1) | 16.8 (14.3 - 21.0) | 10.7 (9.5 - 13.1) | 14.9 (10.3 - 18.6) |
| IgG4 | ChiKaz | 39.9 (31.0 - 46.3) | 35.0 (31.8 - 37.4) | 14.3 (10.1 - 18.8) | 11.3 (9.8 - 13.0) | 10.6 (7.6 - 13.4) |
| IgG4 | ChiKrz | 50.1 (43.3 - 56.3) | 29.9 (27.2 - 33.2) | 9.6 (7.6 - 12.8) | 13.4 (12.1 - 15.6) | 9.5 (6.7 - 13.0) |
| IgG4 | ChiUyg | 44.4 (35.5 - 52.2) | 32.7 (28.2 - 35.9) | 12.0 (8.6 - 15.3) | 11.4 (9.7 - 12.8) | 10.1 (6.4 - 14.9) |
| IgG4 | Croatia | 39.4 (32.3 - 45.9) | 33.2 (29.6 - 35.5) | 12.7 (9.5 - 15.0) | 13.8 (11.3 - 16.9) | 15.1 (11.8 - 17.4) |
| IgG4 | Evenk | 37.8 (31.0 - 45.5) | 31.7 (29.2 - 33.3) | 14.5 (11.1 - 18.0) | 12.7 (11.3 - 14.7) | 15.0 (12.7 - 19.1) |
| IgG4 | Germany | 41.4 (34.6 - 50.3) | 37.4 (32.0 - 39.6) | 11.3 (8.1 - 14.6) | 13.2 (11.0 - 15.2) | 8.9 (6.5 - 11.9) |
| IgG4 | Italy | 40.1 (32.4 - 52.6) | 34.2 (26.8 - 39.3) | 13.4 (7.8 - 17.2) | 13.2 (10.3 - 17.7) | 8.3 (6.3 - 15.6) |
| IgG4 | Kosovo | 37.6 (32.1 - 48.4) | 32.0 (29.3 - 35.2) | 12.5 (9.6 - 16.3) | 13.2 (11.5 - 15.0) | 14.4 (10.6 - 18.2) |
| IgG4 | NewGui | 61.6 (50.6 - 68.3) | 20.1 (17.6 - 24.5) | 7.4 (5.3 - 10.7) | 11.2 (9.1 - 13.3) | 10.8 (8.2 - 14.2) |
| IgG4 | Orkney | 38.0 (29.5 - 45.2) | 32.5 (30.2 - 35.6) | 13.0 (10.1 - 16.7) | 11.9 (10.2 - 14.4) | 15.8 (12.5 - 19.3) |
| IgG4 | Roma | 41.2 (35.5 - 47.4) | 33.0 (30.3 - 35.2) | 13.1 (10.2 - 15.5) | 13.7 (11.6 - 15.8) | 12.8 (9.4 - 16.5) |
| IgG4 | RuKaz | 32.3 (27.4 - 43.3) | 33.3 (30.2 - 36.6) | 16.4 (12.4 - 19.7) | 11.5 (9.7 - 13.3) | 15.4 (12.1 - 18.5) |
| IgG4 | Russia | 34.8 (27.5 - 42.6) | 33.0 (30.4 - 35.6) | 15.4 (11.8 - 18.5) | 11.9 (11.0 - 13.9) | 17.2 (13.2 - 21.0) |
| IgG4 | RuTar | 31.5 (26.2 - 36.8) | 34.0 (31.8 - 35.9) | 16.0 (13.9 - 19.1) | 11.3 (9.7 - 13.0) | 18.1 (14.6 - 21.0) |
| IgG4 | RuYak | 34.2 (29.2 - 40.8) | 31.3 (29.2 - 34.4) | 15.7 (13.1 - 19.2) | 12.4 (10.8 - 14.4) | 16.3 (14.1 - 19.7) |
| IgG4 | SabEng | 51.1 (44.5 - 58.5) | 31.5 (26.9 - 35.9) | 8.2 (5.5 - 10.8) | 15.8 (12.4 - 18.7) | 7.2 (5.5 - 10.1) |
| IgG4 | SabInd | 50.2 (42.7 - 57.0) | 32.0 (28.1 - 34.9) | 8.9 (6.6 - 11.5) | 14.8 (13.0 - 16.7) | 7.5 (5.3 - 10.6) |
| IgG4 | SabJam | 39.3 (33.3 - 46.4) | 35.3 (32.1 - 38.2) | 12.7 (10.8 - 16.0) | 13.2 (11.2 - 15.2) | 11.6 (8.1 - 14.0) |
| IgG4 | Shetland | 37.0 (30.6 - 44.8) | 33.1 (30.8 - 35.4) | 13.1 (11.0 - 15.6) | 11.6 (9.7 - 14.1) | 16.1 (12.8 - 19.2) |
| IgG4 | Sweden | 38.1 (33.1 - 43.4) | 34.0 (31.8 - 36.7) | 13.3 (11.2 - 15.3) | 12.0 (10.2 - 14.1) | 14.0 (11.1 - 17.2) |
| IgG4 | Thailand | 29.5 (23.3 - 35.0) | 34.0 (31.6 - 36.6) | 18.8 (16.1 - 22.0) | 9.6 (8.5 - 11.4) | 16.6 (13.9 - 19.5) |
| IgG4 | TriTob | 35.1 (28.1 - 41.6) | 34.0 (31.5 - 37.2) | 14.9 (11.7 - 18.4) | 11.4 (9.3 - 14.5) | 15.2 (12.5 - 18.6) |
| IgG4 | Turkey | 46.2 (37.8 - 54.5) | 30.4 (27.3 - 33.0) | 10.5 (7.9 - 14.0) | 14.1 (12.5 - 17.6) | 12.8 (9.6 - 15.9) |
| IgG4 | Ugand | 33.1 (26.1 - 42.3) | 31.4 (29.2 - 33.8) | 16.5 (12.7 - 19.0) | 12.0 (9.9 - 14.1) | 18.1 (13.8 - 22.5) |
| IgG4 | TwinsUK | 40.2 (35.2 - 49.7) | 36.4 (32.4 - 38.1) | 12.4 (9.1 - 15.5) | 12.9 (10.5 - 14.8) | 10.5 (6.7 - 14.2) |
| IgG4 | ChiTaj | 36.7 (26.3 - 44.5) | 33.4 (30.9 - 35.5) | 14.8 (11.6 - 20.1) | 11.9 (9.9 - 14.0) | 13.5 (10.1 - 18.3) |

Supplementary Table 10: Health-related Sustainable Development Goals (SDG) indicators used for assessment of country development level[33]. Development indicator and short descriptions are given.

| Health-related SDG indicator | Definition |
|---|---|
| SDG | Overall health-related SDG index (health-related SDG indicators included) |
| MDG | Overall health-related MDX index (Health-related SDG indicators included in the Millennium Development Goals) |
| non-MDG | Health-related SDG indicators not included in the Millennium Development Goals |
| Adolescent birth rate | Birth rates for women aged 10–14 years and women aged 15–19 years, number of livebirths per 1000 women aged 10–14 years and women aged 15–19 years |
| Air pollution mortality | Age-standardised death rate attributable to household air pollution and ambient air pollution, per 100 000 population |
| Alcohol | Risk-weighted prevalence of alcohol consumption, as measured by the SEV for alcohol use, % |
| Disaster | Age-standardised death rate due to exposure to forces of nature, per 100 000 population |
| Family planning need met, modern contraception | Proportion of women of reproductive age (15–49 years) who have their need for family planning satisfied with modern methods, % women aged 15–49 years |
| Hepatitis B | Age-standardised rate of hepatitis B incidence, per 100 000 population |
| HIV | Age-standardised rate of new HIV infections, per 1000 population |
| Household air pollution | Risk-weighted prevalence of household air pollution, as measured by the SEV for household air pollution, % |
| Hygiene | Risk-weighted prevalence of populations with unsafe hygiene (no handwashing with soap), as measured by the SEV for unsafe hygiene, % |
| Intimate partner violence | Age-standardised prevalence of women aged 15 years and older who experienced intimate partner violence, %women aged 15 years and older |
| Malaria | Age-standardised rate of malaria cases, per 1000 population |
| Maternal mortality ratio | Maternal deaths per 100 000 livebirths |
| Mean PM 2.5 | Population-weighted mean levels of PM2.5, $\mu g/m^3$ |
| NCDs | Age-standardised death rate due to cardiovascular disease, cancer, diabetes, and chronic respiratory disease in populations aged 30 - 70 years, per 100 000 population |
| Neglected tropical diseases | Age-standardised prevalence of neglected tropical diseases, per 100 000 population |
| Neonatal mortality | Probability of dying during the first 28 days of life per 1000 livebirths |
| Occupational risk burden | Age-standardised all-cause DALY rate attributable to occupational risks, per 100 000 population |
| Overweight | Prevalence of overweight in children aged 2 - 4 years, % |
| Poisons | Age-standardised death rate due to unintentional poisonings, per 100 000 population |
| Road injuries | Age-standardised death rate due to road traffic injuries, per 100 000 population |
| Sanitation | Risk-weighted prevalence of populations using unsafe or unimproved sanitation, as measured by the SEV for unsafe sanitation, % |
| Skilled birth attendance | Proportion of births attended by skilled health personnel (doctors, nurses, midwives, or country-specific medical staff [e.g., clinical officers]), % |
| Smoking | Age-standardised prevalence of daily smoking in populations aged 10 years and older, % population aged 10 years and older |
| Stunting | Prevalence of stunting in achieving, by 2025, the internationally agreed targets on children under age 5 years, % |
| Suicide | Age-standardised death rate due to self-harm, per 100 000 population |
| Tuberculosis | Age-standardised rate of new and relapsed tuberculosis cases, per 1000 population |
| Under-5 mortality | Probability of dying before age 5 years per 1000 livebirths |
| Universal health coverage tracer | Coverage of universal health coverage tracer interventions for prevention and treatment services, % |
| Violence | Age-standardised death rate due to interpersonal violence, per 100 000 population |
| War | Age-standardised death rate due to collective violence and legal intervention, per 100 000 population |
| WaSH mortality | Age-standardised death rate attributable to unsafe WaSH, per 100 000 population |
| Wasting | Prevalence of wasting in children under age 5 years, % |
| Water | Risk-weighted prevalence of populations using unsafe or unimproved water sources, as measured by the SEV for unsafe water, % |

Supplementary Table 11: United Nation's Human Development Index (HDI) indicators used for assessment of country development level[49]. Development indicator and short descriptions are given.

| HDI indicator | Definition |
|---|---|
| HDI index | Composite measure of three life dimensions: life expectancy, per capita income and education |
| Health index | Life expectancy at birth in 2014; expressed as an index using a minimum value of 20 years and a maximum value of 85 years |
| Life expectancy at birth | Number of years a new-born infant could expect to live if prevailing patterns of age-specific mortality rates at the time of birth stay the same throughout the infant's life |
| Education index | Education index is an average of mean years of schooling (of adults) and expected years of schooling (of children) in 2014, both expressed as an index obtained by scaling with the corresponding maxima |
| GDP | Gross domestic product (GDP) per capita in 2013 |
| Life expectancy - M | Life expectancy at birth, male (years) |
| Life expectancy - F | Life expectancy at birth, female (years) |
| Water (UN) | Accessibility of drinkable tap water |
| Sanitation (UN) | Accessibility of advanced sanitation |

Supplementary Table 12: Development level of 14 countries expressed through 45 indicators. Higher indicator value suggests better conditions impacting on human well-being in a given country. Indicator descriptions are given in Extended Data Table 8 and 9.

| | China | Thailand | Sweden | Germany | England | Scotland | Croatia | Italy | Kosovo | Russia | Turkey | Uganda | Trinidad | Papua |
|---|---|---|---|---|---|---|---|---|---|---|---|---|---|---|
| **SDG index** | 60 | 56 | 85 | 80 | 82 | 82 | 70 | 78 | 65 | 54 | 58 | 31 | 67 | 36 |
| **Disaster** | 39 | 29 | 100 | 100 | 100 | 100 | 100 | 61 | 40 | 57 | 33 | 53 | 100 | 32 |
| **Stunting** | 87 | 85 | 100 | 100 | 100 | 100 | 91 | 100 | 91 | 85 | 86 | 55 | 95 | 51 |
| **Wasting** | 91 | 84 | 100 | 100 | 100 | 100 | 88 | 100 | 87 | 90 | 96 | 82 | 87 | 86 |
| **Overweight** | 62 | 71 | 51 | 47 | 64 | 64 | 46 | 39 | 65 | 46 | 65 | 82 | 81 | 71 |
| **MMR** | 62 | 61 | 79 | 71 | 70 | 70 | 71 | 79 | 68 | 61 | 64 | 28 | 46 | 22 |
| **SBA** | 97 | 99 | 100 | 100 | 99 | 99 | 99 | 99 | 99 | 99 | 94 | 68 | 100 | 65 |
| **Under-5 mort** | 66 | 81 | 94 | 89 | 84 | 84 | 85 | 91 | 75 | 72 | 61 | 32 | 58 | 42 |
| **NN mort** | 69 | 80 | 94 | 91 | 85 | 85 | 84 | 90 | 76 | 75 | 59 | 34 | 51 | 47 |
| **HIV** | 46 | 34 | 65 | 57 | 51 | 51 | 69 | 54 | 57 | 32 | 64 | 15 | 35 | 35 |
| **Tuberculosis** | 45 | 45 | 82 | 86 | 74 | 74 | 69 | 87 | 70 | 41 | 69 | 29 | 62 | 48 |
| **Malaria** | 94 | 26 | 100 | 100 | 100 | 100 | 100 | 100 | 100 | 100 | 100 | 5 | 100 | 6 |
| **Hepatitis B** | 40 | 33 | 85 | 85 | 85 | 85 | 66 | 82 | 66 | 49 | 43 | 33 | 67 | 21 |
| **NTDs** | 89 | 87 | 100 | 100 | 100 | 100 | 100 | 100 | 100 | 99 | 97 | 85 | 98 | 54 |
| **NCDs** | 58 | 61 | 87 | 77 | 78 | 78 | 62 | 85 | 54 | 41 | 73 | 46 | 47 | 19 |
| **Suicide** | 59 | 42 | 51 | 55 | 64 | 64 | 47 | 71 | 44 | 21 | 81 | 43 | 46 | 42 |
| **Alcohol** | 74 | 74 | 57 | 54 | 57 | 57 | 54 | 68 | 59 | 7 | 90 | 58 | 65 | 87 |
| **Road injuries** | 49 | 36 | 94 | 88 | 94 | 94 | 72 | 75 | 69 | 53 | 66 | 35 | 55 | 37 |
| **FP need met** | 86 | 66 | 91 | 92 | 95 | 95 | 34 | 86 | 37 | 75 | 70 | 41 | 76 | 47 |
| **Adol birth rate** | 69 | 54 | 84 | 86 | 73 | 73 | 80 | 87 | 70 | 67 | 61 | 24 | 60 | 44 |
| **UHC Tracer** | 81 | 69 | 99 | 96 | 100 | 100 | 84 | 97 | 84 | 85 | 82 | 69 | 93 | 45 |
| **Air poll mort** | 48 | 61 | 96 | 83 | 83 | 83 | 68 | 83 | 58 | 62 | 71 | 33 | 74 | 28 |
| **WaSH mort** | 75 | 49 | 87 | 84 | 77 | 77 | 88 | 96 | 88 | 79 | 76 | 22 | 64 | 21 |
| **Poisons** | 50 | 69 | 71 | 94 | 75 | 75 | 84 | 81 | 73 | 47 | 80 | 37 | 71 | 44 |
| **Smoking** | 52 | 60 | 75 | 46 | 55 | 55 | 32 | 52 | 43 | 41 | 47 | 87 | 68 | 36 |
| **IPV** | 96 | 57 | 80 | 70 | 80 | 80 | 91 | 66 | 82 | 64 | 53 | 18 | 73 | 69 |
| **Water** | 500 | 34 | 100 | 100 | 100 | 100 | 81 | 100 | 74 | 86 | 29 | 26 | 47 | 14 |
| **Sanitation** | 66 | 98 | 100 | 100 | 100 | 100 | 97 | 100 | 85 | 79 | 87 | 20 | 90 | 20 |
| **Hygiene** | 38 | 46 | 95 | 89 | 93 | 93 | 43 | 86 | 43 | 69 | 28 | 3 | 53 | 16 |
| **HH air poll** | 79 | 88 | 100 | 100 | 100 | 100 | 94 | 100 | 78 | 98 | 98 | 2 | 100 | 55 |
| **Occ risk burden** | 43 | 54 | 81 | 71 | 74 | 74 | 63 | 72 | 60 | 67 | 57 | 31 | 73 | 9 |
| **Mean PM 2.5** | 25 | 46 | 81 | 62 | 65 | 65 | 51 | 53 | 51 | 58 | 37 | 24 | 62 | 62 |
| **Violence** | 69 | 34 | 74 | 81 | 86 | 86 | 72 | 77 | 55 | 25 | 58 | 39 | 19 | 36 |
| **War** | 100 | 100 | 100 | 100 | 100 | 100 | 100 | 100 | 100 | 31 | 19 | 100 | 100 | 100 |
| **MDG index** | 70 | 63 | 94 | 92 | 90 | 90 | 80 | 92 | 76 | 75 | 67 | 33 | 69 | 33 |
| **Non-MDG index** | 55 | 54 | 80 | 73 | 78 | 78 | 64 | 70 | 60 | 46 | 54 | 29 | 67 | 37 |
| **HDI index** | 73 | 73 | 91 | 92 | 91 | 91 | 82 | 87 | 77 | 80 | 76 | 48 | 77 | 51 |
| **Life exp** | 76 | 74 | 82 | 81 | 81 | 81 | 77 | 83 | 75 | 70 | 75 | 59 | 70 | 63 |

| | | | | | | | | | | | | | | |
|---|---|---|---|---|---|---|---|---|---|---|---|---|---|---|
| **Life exp F** | 78 | 78 | 84 | 83 | 83 | 83 | 81 | 86 | 78 | 76 | 79 | 61 | 74 | 65 |
| **Life exp M** | 75 | 71 | 81 | 79 | 79 | 79 | 74 | 81 | 72 | 65 | 72 | 57 | 67 | 61 |
| **GDP** | 11525 | 13932 | 43741 | 43207 | 37017 | 37017 | 20063 | 34167 | 12893 | 23564 | 18660 | 1368 | 29469 | 2458 |
| **Education index** | 61 | 62 | 84 | 89 | 89 | 89 | 78 | 78 | 75 | 81 | 66 | 45 | 71 | 41 |
| **Health Index** | 86 | 84 | 96 | 94 | 93 | 93 | 88 | 97 | 85 | 77 | 85 | 59 | 78 | 66 |
| **Water (UN)** | 96 | 98 | 100 | 100 | 100 | 100 | 100 | 100 | 99 | 97 | 100 | 79 | 95 | 40 |
| **Sanitation (UN)** | 77 | 93 | 99 | 99 | 99 | 99 | 97 | 100 | 96 | 72 | 95 | 19 | 92 | 19 |

*MMR = maternal mortality ratio; SBA = skilled birth attendance; Under-5 mort = Under-5 mortality; NN mort = neonatal mortality; NTDs = neglected tropical diseases; NCDs = non-communicable diseases; FP need met = family planning need met, modern contraception; Adol birth rate = adolescent birth rate; UHC = universal health coverage; Air poll mort = air pollution mortality; WaSH Mort = water, sanitation, and hygiene mortality; IPV = intimate partner violence; HH air poll = household air pollution; MDG index = Millennium Development Goals Index; non-MDG index = health-related Sustainable Development Goals not included in MDG; HDI = Human Development Index; Life exp = Life expectancy at birth; Life exp F = Life expectancy – F; Life exp M = Life expectancy - M

Supplementary Table 13: Correlations of IgG Fc derived glycan traits with participant’s country of residence development indicators. Pearson's correlation coefficient with accompanying P values are given (n=14). Abbreviations: MMR = maternal mortality ratio; SBA = skilled birth attendance; Nnmort = neonatal mortality; NTDs = neglected tropical diseases; NCDs = non-communicable diseases; FPneedmet = family planning need met, modern contraception; Adol = Adolescent birth rate; UHC = universal health coverage; Air poll mort = Air pollution mortality; WaSH = water, sanitation, and hygiene; IPV = intimate partner violence; HHairpoll = household air pollution; MDG = Millenium Development Goals; HDI = Human development Index; SDG = Sustainable Development Goals. Development indicators are described in Etended Data Tables 8 and 9.

| Glycan Trait | Development index/indicator | r | P value | adjusted P value |
|---|---|---|---|---|
| IgG1_Monogalactosylation | MDG | 0.969434756 | 1.10E-08 | 7.44E-06 |
| IgG1_Monogalactosylation | Stunting | 0.967055014 | 1.72E-08 | 1.16E-05 |
| IgG1_Monogalactosylation | HDI | 0.965825352 | 2.14E-08 | 1.44E-05 |
| IgG1_Agalactosylation | UHCTracer | -0.95037426 | 1.94E-07 | 1.31E-04 |
| IgG1_Agalactosylation | OccRiskBurden | -0.950003444 | 2.02E-07 | 1.37E-04 |
| IgG1_Monogalactosylation | UHCTracer | 0.945363091 | 3.41E-07 | 2.30E-04 |
| IgG1_Agalactosylation | Stunting | -0.9420058 | 4.84E-07 | 3.27E-04 |
| IgG1_Monogalactosylation | SDG | 0.940338227 | 5.72E-07 | 3.86E-04 |
| IgG1_Monogalactosylation | EduIndx_2014 | 0.937368449 | 7.61E-07 | 5.14E-04 |
| IgG1_Monogalactosylation | OccRiskBurden | 0.93089952 | 1.35E-06 | 9.13E-04 |
| IgG1_Digalactosylation | SBA | 0.930467933 | 1.40E-06 | 9.47E-04 |
| IgG1_Digalactosylation | Stunting | 0.923405123 | 2.47E-06 | 1.67E-03 |
| IgG1_Monogalactosylation | LifeExp_F | 0.916985412 | 3.94E-06 | 2.66E-03 |
| IgG1_Monogalactosylation | AirPollMort | 0.914275509 | 4.75E-06 | 3.21E-03 |
| IgG1_Agalactosylation | HDI | -0.906023161 | 8.10E-06 | 5.47E-03 |
| IgG1_Monogalactosylation | HealthIndx_2014 | 0.901032539 | 1.09E-05 | 7.38E-03 |
| IgG1_Monogalactosylation | LifeExp | 0.901010147 | 1.09E-05 | 7.39E-03 |
| IgG1_Agalactosylation | MDG | -0.899285831 | 1.21E-05 | 8.16E-03 |
| IgG1_Monogalactosylation | Non.MDG | 0.898094676 | 1.29E-05 | 8.73E-03 |
| IgG1_Agalactosylation | EduIndx_2014 | -0.895317502 | 1.51E-05 | 1.02E-02 |
| IgG1_Monogalactosylation | HepatB | 0.895249048 | 1.52E-05 | 1.02E-02 |
| IgG1_Agalactosylation | Water_UN | -0.891382686 | 1.87E-05 | 1.26E-02 |
| IgG2_Monogalactosylation | Stunting | 0.890858888 | 1.92E-05 | 1.30E-02 |
| IgG1_Agalactosylation | SBA | -0.890488563 | 1.96E-05 | 1.32E-02 |
| IgG1_Monogalactosylation | MMR | 0.890356305 | 1.97E-05 | 1.33E-02 |
| IgG1_Monogalactosylation | GDP_2013 | 0.888528777 | 2.17E-05 | 1.46E-02 |
| IgG1_Monogalactosylation | Sanitation_UN | 0.884836577 | 2.62E-05 | 1.77E-02 |
| IgG1_Monogalactosylation | Hygiene | 0.881989588 | 3.01E-05 | 2.03E-02 |
| IgG1_Digalactosylation | Water_UN | 0.880105424 | 3.30E-05 | 2.22E-02 |
| IgG1_Monogalactosylation | SBA | 0.875839535 | 4.03E-05 | 2.72E-02 |
| IgG1_Monogalactosylation | WaSHMort | 0.87368469 | 4.44E-05 | 3.00E-02 |
| IgG4_Monogalactosylation | Water_UN | 0.872607839 | 4.66E-05 | 3.15E-02 |
| IgG1_Agalactosylation | AirPollMort | -0.871570709 | 4.88E-05 | 3.30E-02 |
| IgG1_Agalactosylation | SDG | -0.871295686 | 4.94E-05 | 3.34E-02 |
| IgG1_Monogalactosylation | AdolBirthRate | 0.870585663 | 5.10E-05 | 3.44E-02 |
| IgG1_Monogalactosylation | LifeExp_M | 0.868743447 | 5.53E-05 | 3.73E-02 |
| IgG1_Digalactosylation | Sanitation_UN | 0.867063308 | 5.95E-05 | 4.01E-02 |
| IgG1_Digalactosylation | OccRiskBurden | 0.866441892 | 6.11E-05 | 4.12E-02 |
| IgG1_Monogalactosylation | Under5Mort | 0.866368769 | 6.13E-05 | 4.14E-02 |
| IgG2_Monogalactosylation | HDI | 0.864097812 | 6.74E-05 | 4.55E-02 |
| IgG2_Monogalactosylation | OccRiskBurden | 0.864088251 | 6.75E-05 | 4.55E-02 |
| IgG2_Monogalactosylation | MDG | 0.863881556 | 6.80E-05 | 4.59E-02 |
| IgG2_Monogalactosylation | AirPollMort | 0.862569569 | 7.19E-05 | 4.85E-02 |
| IgG1_Agalactosylation | Sanitation_UN | -0.861935177 | 7.38E-05 | 4.98E-02 |
| IgG1_Digalactosylation | MDG | 0.861366104 | 7.55E-05 | 5.10E-02 |
| IgG1_Digalactosylation | HDI | 0.860780423 | 7.74E-05 | 5.22E-02 |
| IgG2_Monogalactosylation | UHCTracer | 0.853575289 | 1.03E-04 | 6.95E-02 |
| IgG1_Digalactosylation | UHCTracer | 0.848991406 | 1.23E-04 | 8.28E-02 |
| IgG4_Monogalactosylation | OccRiskBurden | 0.846935798 | 1.32E-04 | 8.94E-02 |
| IgG4_Monogalactosylation | UHCTracer | 0.845494245 | 1.40E-04 | 9.42E-02 |
| IgG2_Monogalactosylation | SDG | 0.844836201 | 1.43E-04 | 9.65E-02 |
| IgG2_Agalactosylation | Water_UN | -0.843662929 | 1.49E-04 | 1.01E-01 |
| IgG4_Monogalactosylation | Stunting | 0.842223049 | 1.57E-04 | 1.06E-01 |
| IgG2_Monogalactosylation | LifeExp_F | 0.84175077 | 1.60E-04 | 1.08E-01 |
| IgG1_Digalactosylation | LifeExp_F | 0.840123908 | 1.69E-04 | 1.14E-01 |
| IgG1_Agalactosylation | Non.MDG | -0.839575445 | 1.73E-04 | 1.16E-01 |
| IgG2_Monogalactosylation | GDP_2013 | 0.837973778 | 1.82E-04 | 1.23E-01 |
| IgG1_Agalactosylation | HepatB | -0.833457052 | 2.13E-04 | 1.44E-01 |

| | | | | |
|---|---|---|---|---|
| IgG1_Digalactosylation | MMR | 0.829385853 | 2.44E-04 | 1.65E-01 |
| IgG4_Monogalactosylation | SBA | 0.827296068 | 2.61E-04 | 1.76E-01 |
| IgG1_Monogalactosylation | Water_UN | 0.826515112 | 2.68E-04 | 1.81E-01 |
| IgG2_Agalactosylation | Stunting | -0.826331354 | 2.69E-04 | 1.82E-01 |
| IgG1_Agalactosylation | LifeExp_F | -0.824494857 | 2.86E-04 | 1.93E-01 |
| IgG2_Agalactosylation | SBA | -0.823894008 | 2.91E-04 | 1.97E-01 |
| IgG2_Monogalactosylation | Hygiene | 0.823431265 | 2.96E-04 | 2.00E-01 |
| IgG1_Monogalactosylation | RoadInjuries | 0.823239508 | 2.97E-04 | 2.01E-01 |
| IgG1_Monogalactosylation | Nnmort | 0.822245754 | 3.07E-04 | 2.07E-01 |
| IgG1_Agalactosylation | GDP_2013 | -0.821517563 | 3.14E-04 | 2.12E-01 |
| IgG1_Monogalactosylation | NCDs | 0.821496789 | 3.14E-04 | 2.12E-01 |
| IgG1_Agalactosylation | MMR | -0.821489913 | 3.14E-04 | 2.12E-01 |
| IgG2_Monogalactosylation | LifeExp | 0.820466756 | 3.24E-04 | 2.19E-01 |
| IgG2_Monogalactosylation | HealthIndx_2014 | 0.820204125 | 3.27E-04 | 2.21E-01 |
| IgG2_Monogalactosylation | Non.MDG | 0.812966233 | 4.08E-04 | 2.75E-01 |
| IgG2_Monogalactosylation | SBA | 0.812231749 | 4.17E-04 | 2.81E-01 |
| IgG4_Agalactosylation | Water_UN | -0.809870949 | 4.47E-04 | 3.01E-01 |
| IgG2_Agalactosylation | OccRiskBurden | -0.809791983 | 4.48E-04 | 3.02E-01 |
| IgG2_Monogalactosylation | Sanitation_UN | 0.807061294 | 4.85E-04 | 3.27E-01 |
| IgG1_Digalactosylation | LifeExp | 0.806219491 | 4.96E-04 | 3.35E-01 |
| IgG1_Agalactosylation | Hygiene | -0.806160418 | 4.97E-04 | 3.36E-01 |
| IgG1_Digalactosylation | HealthIndx_2014 | 0.806030521 | 4.99E-04 | 3.37E-01 |
| IgG1_Digalactosylation | SDG | 0.804797471 | 5.17E-04 | 3.49E-01 |
| IgG1_Digalactosylation | EduIndx_2014 | 0.802333078 | 5.54E-04 | 3.74E-01 |
| IgG1_Agalactosylation | WaSHMort | -0.798814855 | 6.11E-04 | 4.12E-01 |
| IgG1_Agalactosylation | HealthIndx_2014 | -0.797763059 | 6.29E-04 | 4.25E-01 |
| IgG1_Agalactosylation | LifeExp | -0.797752927 | 6.29E-04 | 4.25E-01 |
| IgG2_Monogalactosylation | EduIndx_2014 | 0.79697559 | 6.43E-04 | 4.34E-01 |
| IgG1_Digalactosylation | WaSHMort | 0.796649312 | 6.48E-04 | 4.38E-01 |
| IgG2_Monogalactosylation | Under5Mort | 0.794179045 | 6.93E-04 | 4.68E-01 |
| IgG2_Digalactosylation | SBA | 0.793871394 | 6.99E-04 | 4.72E-01 |
| IgG1_Bisecting | Water_UN | 0.791100782 | 7.52E-04 | 5.08E-01 |
| IgG1_Bisecting | WaSHMort | 0.789357384 | 7.87E-04 | 5.32E-01 |
| IgG2_Monogalactosylation | LifeExp_M | 0.787301248 | 8.31E-04 | 5.61E-01 |
| IgG1_Monogalactosylation | HHAirPoll | 0.786664858 | 8.44E-04 | 5.70E-01 |
| IgG1_Digalactosylation | Under5Mort | 0.786047351 | 8.58E-04 | 5.79E-01 |
| IgG2_Monogalactosylation | MMR | 0.785717063 | 8.65E-04 | 5.84E-01 |
| IgG1_Digalactosylation | AirPollMort | 0.785520456 | 8.70E-04 | 5.87E-01 |
| IgG2_Agalactosylation | Sanitation_UN | -0.783580847 | 9.14E-04 | 6.17E-01 |
| IgG2_Monogalactosylation | HepatB | 0.780914085 | 9.77E-04 | 6.60E-01 |
| IgG1_Agalactosylation | Under5Mort | -0.779220205 | 1.02E-03 | 6.88E-01 |
| IgG2_Agalactosylation | UHCTracer | -0.77700753 | 1.08E-03 | 7.27E-01 |
| IgG4_Monogalactosylation | HDI | 0.776086511 | 1.10E-03 | 7.43E-01 |
| IgG2_Monogalactosylation | AdolBirthRate | 0.774879532 | 1.13E-03 | 7.66E-01 |
| IgG4_Monogalactosylation | MDG | 0.771267904 | 1.24E-03 | 8.35E-01 |
| IgG1_Bisecting | MMR | 0.770597406 | 1.26E-03 | 8.49E-01 |
| IgG1_Monogalactosylation | Wasting | 0.767918251 | 1.34E-03 | 9.04E-01 |
| IgG4_Monogalactosylation | Sanitation_UN | 0.767152449 | 1.36E-03 | 9.21E-01 |
| IgG2_Monogalactosylation | NCDs | 0.766742835 | 1.38E-03 | 9.29E-01 |
| IgG4_Monogalactosylation | EduIndx_2014 | 0.764681027 | 1.44E-03 | 9.75E-01 |
| IgG1_Sialylation | HDI | -0.512835432 | 6.08E-02 | 1.00E+00 |
| IgG1_Bisecting | HDI | 0.647646104 | 1.23E-02 | 1.00E+00 |
| IgG2_Agalactosylation | HDI | -0.756061358 | 1.76E-03 | 1.00E+00 |
| IgG2_Digalactosylation | HDI | 0.649999603 | 1.19E-02 | 1.00E+00 |
| IgG2_Sialylation | HDI | 0.121510885 | 6.79E-01 | 1.00E+00 |
| IgG2_Bisecting | HDI | 0.529085703 | 5.17E-02 | 1.00E+00 |
| IgG4_Agalactosylation | HDI | -0.395134154 | 1.62E-01 | 1.00E+00 |
| IgG4_Digalactosylation | HDI | 0.240297289 | 4.08E-01 | 1.00E+00 |
| IgG4_Sialylation | HDI | -0.052923302 | 8.57E-01 | 1.00E+00 |
| IgG4_Bisecting | HDI | 0.030439221 | 9.18E-01 | 1.00E+00 |
| IgG1_Sialylation | SDG | -0.468841347 | 9.08E-02 | 1.00E+00 |
| IgG1_Bisecting | SDG | 0.529506192 | 5.15E-02 | 1.00E+00 |
| IgG2_Agalactosylation | SDG | -0.704767262 | 4.88E-03 | 1.00E+00 |
| IgG2_Digalactosylation | SDG | 0.58127416 | 2.92E-02 | 1.00E+00 |
| IgG2_Sialylation | SDG | 0.073926851 | 8.02E-01 | 1.00E+00 |
| IgG2_Bisecting | SDG | 0.411231552 | 1.44E-01 | 1.00E+00 |
| IgG4_Agalactosylation | SDG | -0.316231061 | 2.71E-01 | 1.00E+00 |
| IgG4_Monogalactosylation | SDG | 0.725112788 | 3.34E-03 | 1.00E+00 |
| IgG4_Digalactosylation | SDG | 0.16147417 | 5.81E-01 | 1.00E+00 |
| IgG4_Sialylation | SDG | -0.112824422 | 7.01E-01 | 1.00E+00 |
| IgG4_Bisecting | SDG | 0.009620853 | 9.74E-01 | 1.00E+00 |
| IgG1_Agalactosylation | Disaster | -0.68565858 | 6.79E-03 | 1.00E+00 |
| IgG1_Monogalactosylation | Disaster | 0.6608941 | 1.01E-02 | 1.00E+00 |

| | | | | |
|---|---|---|---|---|
| IgG1_Digalactosylation | Disaster | 0.479713009 | 8.26E-02 | 1.00E+00 |
| IgG1_Sialylation | Disaster | 0.111545691 | 7.04E-01 | 1.00E+00 |
| IgG1_Bisecting | Disaster | 0.209030447 | 4.73E-01 | 1.00E+00 |
| IgG2_Agalactosylation | Disaster | -0.411580883 | 1.44E-01 | 1.00E+00 |
| IgG2_Monogalactosylation | Disaster | 0.587857022 | 2.70E-02 | 1.00E+00 |
| IgG2_Digalactosylation | Disaster | 0.188855976 | 5.18E-01 | 1.00E+00 |
| IgG2_Sialylation | Disaster | 0.157024067 | 5.92E-01 | 1.00E+00 |
| IgG2_Bisecting | Disaster | -0.000407641 | 9.99E-01 | 1.00E+00 |
| IgG4_Agalactosylation | Disaster | -0.285038409 | 3.23E-01 | 1.00E+00 |
| IgG4_Monogalactosylation | Disaster | 0.582205084 | 2.89E-02 | 1.00E+00 |
| IgG4_Digalactosylation | Disaster | 0.087805905 | 7.65E-01 | 1.00E+00 |
| IgG4_Sialylation | Disaster | 0.078446819 | 7.90E-01 | 1.00E+00 |
| IgG4_Bisecting | Disaster | -0.013813054 | 9.63E-01 | 1.00E+00 |
| IgG1_Sialylation | Stunting | -0.518666635 | 5.74E-02 | 1.00E+00 |
| IgG1_Bisecting | Stunting | 0.640431247 | 1.36E-02 | 1.00E+00 |
| IgG2_Digalactosylation | Stunting | 0.737032466 | 2.63E-03 | 1.00E+00 |
| IgG2_Sialylation | Stunting | 0.191858217 | 5.11E-01 | 1.00E+00 |
| IgG2_Bisecting | Stunting | 0.485180471 | 7.87E-02 | 1.00E+00 |
| IgG4_Agalactosylation | Stunting | -0.524210355 | 5.43E-02 | 1.00E+00 |
| IgG4_Digalactosylation | Stunting | 0.390887956 | 1.67E-01 | 1.00E+00 |
| IgG4_Sialylation | Stunting | 0.039326919 | 8.94E-01 | 1.00E+00 |
| IgG4_Bisecting | Stunting | -0.077366962 | 7.93E-01 | 1.00E+00 |
| IgG1_Agalactosylation | Wasting | -0.622517725 | 1.74E-02 | 1.00E+00 |
| IgG1_Digalactosylation | Wasting | 0.558535171 | 3.79E-02 | 1.00E+00 |
| IgG1_Sialylation | Wasting | -0.54822426 | 4.24E-02 | 1.00E+00 |
| IgG1_Bisecting | Wasting | 0.425687396 | 1.29E-01 | 1.00E+00 |
| IgG2_Agalactosylation | Wasting | -0.486896407 | 7.74E-02 | 1.00E+00 |
| IgG2_Monogalactosylation | Wasting | 0.680690058 | 7.37E-03 | 1.00E+00 |
| IgG2_Digalactosylation | Wasting | 0.376317069 | 1.85E-01 | 1.00E+00 |
| IgG2_Sialylation | Wasting | -0.160863862 | 5.83E-01 | 1.00E+00 |
| IgG2_Bisecting | Wasting | 0.427929189 | 1.27E-01 | 1.00E+00 |
| IgG4_Agalactosylation | Wasting | 0.004851465 | 9.87E-01 | 1.00E+00 |
| IgG4_Monogalactosylation | Wasting | 0.475830488 | 8.55E-02 | 1.00E+00 |
| IgG4_Digalactosylation | Wasting | -0.15516085 | 5.96E-01 | 1.00E+00 |
| IgG4_Sialylation | Wasting | -0.375822845 | 1.85E-01 | 1.00E+00 |
| IgG4_Bisecting | Wasting | 0.259264679 | 3.71E-01 | 1.00E+00 |
| IgG1_Agalactosylation | Overweight | 0.440063315 | 1.15E-01 | 1.00E+00 |
| IgG1_Monogalactosylation | Overweight | -0.597639054 | 2.40E-02 | 1.00E+00 |
| IgG1_Digalactosylation | Overweight | -0.474084685 | 8.68E-02 | 1.00E+00 |
| IgG1_Sialylation | Overweight | 0.651490247 | 1.16E-02 | 1.00E+00 |
| IgG1_Bisecting | Overweight | -0.540272828 | 4.61E-02 | 1.00E+00 |
| IgG2_Agalactosylation | Overweight | 0.38036389 | 1.80E-01 | 1.00E+00 |
| IgG2_Monogalactosylation | Overweight | -0.543930227 | 4.44E-02 | 1.00E+00 |
| IgG2_Digalactosylation | Overweight | -0.383445469 | 1.76E-01 | 1.00E+00 |
| IgG2_Sialylation | Overweight | 0.296425717 | 3.03E-01 | 1.00E+00 |
| IgG2_Bisecting | Overweight | -0.586556576 | 2.75E-02 | 1.00E+00 |
| IgG4_Agalactosylation | Overweight | 0.007801878 | 9.79E-01 | 1.00E+00 |
| IgG4_Monogalactosylation | Overweight | -0.38011317 | 1.80E-01 | 1.00E+00 |
| IgG4_Digalactosylation | Overweight | 0.081318441 | 7.82E-01 | 1.00E+00 |
| IgG4_Sialylation | Overweight | 0.301473685 | 2.95E-01 | 1.00E+00 |
| IgG4_Bisecting | Overweight | -0.414746144 | 1.40E-01 | 1.00E+00 |
| IgG1_Sialylation | MMR | -0.626475779 | 1.65E-02 | 1.00E+00 |
| IgG2_Agalactosylation | MMR | -0.743594726 | 2.30E-03 | 1.00E+00 |
| IgG2_Digalactosylation | MMR | 0.676905841 | 7.84E-03 | 1.00E+00 |
| IgG2_Sialylation | MMR | 0.176345168 | 5.46E-01 | 1.00E+00 |
| IgG2_Bisecting | MMR | 0.698558966 | 5.45E-03 | 1.00E+00 |
| IgG4_Agalactosylation | MMR | -0.426005718 | 1.29E-01 | 1.00E+00 |
| IgG4_Monogalactosylation | MMR | 0.707747908 | 4.63E-03 | 1.00E+00 |
| IgG4_Digalactosylation | MMR | 0.315794855 | 2.71E-01 | 1.00E+00 |
| IgG4_Sialylation | MMR | 0.003984067 | 9.89E-01 | 1.00E+00 |
| IgG4_Bisecting | MMR | 0.142564154 | 6.27E-01 | 1.00E+00 |
| IgG1_Sialylation | SBA | -0.506135755 | 6.48E-02 | 1.00E+00 |
| IgG1_Bisecting | SBA | 0.676067183 | 7.94E-03 | 1.00E+00 |
| IgG2_Sialylation | SBA | 0.260388753 | 3.69E-01 | 1.00E+00 |
| IgG2_Bisecting | SBA | 0.507473706 | 6.40E-02 | 1.00E+00 |
| IgG4_Agalactosylation | SBA | -0.627201116 | 1.64E-02 | 1.00E+00 |
| IgG4_Digalactosylation | SBA | 0.540164578 | 4.61E-02 | 1.00E+00 |
| IgG4_Sialylation | SBA | 0.171052148 | 5.59E-01 | 1.00E+00 |
| IgG4_Bisecting | SBA | -0.188359717 | 5.19E-01 | 1.00E+00 |
| IgG1_Sialylation | Under5Mort | -0.586299164 | 2.76E-02 | 1.00E+00 |
| IgG1_Bisecting | Under5Mort | 0.589730559 | 2.64E-02 | 1.00E+00 |
| IgG2_Agalactosylation | Under5Mort | -0.712602803 | 4.23E-03 | 1.00E+00 |
| IgG2_Digalactosylation | Under5Mort | 0.656419002 | 1.08E-02 | 1.00E+00 |

| | | | | |
|---|---|---|---|---|
| IgG2_Sialylation | Under5Mort | 0.095756562 | 7.45E-01 | 1.00E+00 |
| IgG2_Bisecting | Under5Mort | 0.534202452 | 4.91E-02 | 1.00E+00 |
| IgG4_Agalactosylation | Under5Mort | -0.318690201 | 2.67E-01 | 1.00E+00 |
| IgG4_Monogalactosylation | Under5Mort | 0.66574538 | 9.35E-03 | 1.00E+00 |
| IgG4_Digalactosylation | Under5Mort | 0.219735454 | 4.50E-01 | 1.00E+00 |
| IgG4_Sialylation | Under5Mort | -0.114031357 | 6.98E-01 | 1.00E+00 |
| IgG4_Bisecting | Under5Mort | 0.013005203 | 9.65E-01 | 1.00E+00 |
| IgG1_Agalactosylation | Nnmort | -0.715323835 | 4.03E-03 | 1.00E+00 |
| IgG1_Digalactosylation | Nnmort | 0.727820535 | 3.17E-03 | 1.00E+00 |
| IgG1_Sialylation | Nnmort | -0.606915915 | 2.14E-02 | 1.00E+00 |
| IgG1_Bisecting | Nnmort | 0.558862093 | 3.78E-02 | 1.00E+00 |
| IgG2_Agalactosylation | Nnmort | -0.646188088 | 1.25E-02 | 1.00E+00 |
| IgG2_Monogalactosylation | Nnmort | 0.732182185 | 2.91E-03 | 1.00E+00 |
| IgG2_Digalactosylation | Nnmort | 0.602425825 | 2.26E-02 | 1.00E+00 |
| IgG2_Sialylation | Nnmort | 0.054118823 | 8.54E-01 | 1.00E+00 |
| IgG2_Bisecting | Nnmort | 0.533005119 | 4.97E-02 | 1.00E+00 |
| IgG4_Agalactosylation | Nnmort | -0.229448008 | 4.30E-01 | 1.00E+00 |
| IgG4_Monogalactosylation | Nnmort | 0.600980357 | 2.30E-02 | 1.00E+00 |
| IgG4_Digalactosylation | Nnmort | 0.137171605 | 6.40E-01 | 1.00E+00 |
| IgG4_Sialylation | Nnmort | -0.187733241 | 5.20E-01 | 1.00E+00 |
| IgG4_Bisecting | Nnmort | 0.056595872 | 8.48E-01 | 1.00E+00 |
| IgG1_Agalactosylation | HIV | -0.461388721 | 9.68E-02 | 1.00E+00 |
| IgG1_Monogalactosylation | HIV | 0.599964338 | 2.33E-02 | 1.00E+00 |
| IgG1_Digalactosylation | HIV | 0.418003706 | 1.37E-01 | 1.00E+00 |
| IgG1_Sialylation | HIV | -0.454826608 | 1.02E-01 | 1.00E+00 |
| IgG1_Bisecting | HIV | 0.688790351 | 6.44E-03 | 1.00E+00 |
| IgG2_Agalactosylation | HIV | -0.275785121 | 3.40E-01 | 1.00E+00 |
| IgG2_Monogalactosylation | HIV | 0.439740809 | 1.16E-01 | 1.00E+00 |
| IgG2_Digalactosylation | HIV | 0.194577069 | 5.05E-01 | 1.00E+00 |
| IgG2_Sialylation | HIV | -0.157795771 | 5.90E-01 | 1.00E+00 |
| IgG2_Bisecting | HIV | 0.662516552 | 9.83E-03 | 1.00E+00 |
| IgG4_Agalactosylation | HIV | 0.036314938 | 9.02E-01 | 1.00E+00 |
| IgG4_Monogalactosylation | HIV | 0.318221483 | 2.68E-01 | 1.00E+00 |
| IgG4_Digalactosylation | HIV | -0.170423567 | 5.60E-01 | 1.00E+00 |
| IgG4_Sialylation | HIV | -0.182679967 | 5.32E-01 | 1.00E+00 |
| IgG4_Bisecting | HIV | 0.401165555 | 1.55E-01 | 1.00E+00 |
| IgG1_Agalactosylation | Tuberculosis | -0.603069247 | 2.24E-02 | 1.00E+00 |
| IgG1_Monogalactosylation | Tuberculosis | 0.76203146 | 1.54E-03 | 1.00E+00 |
| IgG1_Digalactosylation | Tuberculosis | 0.517171826 | 5.82E-02 | 1.00E+00 |
| IgG1_Sialylation | Tuberculosis | -0.490455499 | 7.50E-02 | 1.00E+00 |
| IgG1_Bisecting | Tuberculosis | 0.524246562 | 5.43E-02 | 1.00E+00 |
| IgG2_Agalactosylation | Tuberculosis | -0.442737669 | 1.13E-01 | 1.00E+00 |
| IgG2_Monogalactosylation | Tuberculosis | 0.665653408 | 9.36E-03 | 1.00E+00 |
| IgG2_Digalactosylation | Tuberculosis | 0.310149244 | 2.81E-01 | 1.00E+00 |
| IgG2_Sialylation | Tuberculosis | -0.167224 | 5.68E-01 | 1.00E+00 |
| IgG2_Bisecting | Tuberculosis | 0.493822105 | 7.27E-02 | 1.00E+00 |
| IgG4_Agalactosylation | Tuberculosis | -0.004363498 | 9.88E-01 | 1.00E+00 |
| IgG4_Monogalactosylation | Tuberculosis | 0.484600794 | 7.91E-02 | 1.00E+00 |
| IgG4_Digalactosylation | Tuberculosis | -0.169672041 | 5.62E-01 | 1.00E+00 |
| IgG4_Sialylation | Tuberculosis | -0.30977615 | 2.81E-01 | 1.00E+00 |
| IgG4_Bisecting | Tuberculosis | 0.308524401 | 2.83E-01 | 1.00E+00 |
| IgG1_Agalactosylation | Malaria | -0.107121449 | 7.15E-01 | 1.00E+00 |
| IgG1_Monogalactosylation | Malaria | 0.147136165 | 6.16E-01 | 1.00E+00 |
| IgG1_Digalactosylation | Malaria | 0.089636204 | 7.61E-01 | 1.00E+00 |
| IgG1_Sialylation | Malaria | -0.125945371 | 6.68E-01 | 1.00E+00 |
| IgG1_Bisecting | Malaria | 0.415959578 | 1.39E-01 | 1.00E+00 |
| IgG2_Agalactosylation | Malaria | 0.01787872 | 9.52E-01 | 1.00E+00 |
| IgG2_Monogalactosylation | Malaria | -0.059131692 | 8.41E-01 | 1.00E+00 |
| IgG2_Digalactosylation | Malaria | -0.029620599 | 9.20E-01 | 1.00E+00 |
| IgG2_Sialylation | Malaria | 0.135425653 | 6.44E-01 | 1.00E+00 |
| IgG2_Bisecting | Malaria | 0.411093709 | 1.44E-01 | 1.00E+00 |
| IgG4_Agalactosylation | Malaria | -0.053910439 | 8.55E-01 | 1.00E+00 |
| IgG4_Monogalactosylation | Malaria | 0.069863161 | 8.12E-01 | 1.00E+00 |
| IgG4_Digalactosylation | Malaria | -0.002788222 | 9.92E-01 | 1.00E+00 |
| IgG4_Sialylation | Malaria | 0.097345335 | 7.41E-01 | 1.00E+00 |
| IgG4_Bisecting | Malaria | 0.227295177 | 4.35E-01 | 1.00E+00 |
| IgG1_Digalactosylation | HepatB | 0.690722326 | 6.24E-03 | 1.00E+00 |
| IgG1_Sialylation | HepatB | -0.313363699 | 2.75E-01 | 1.00E+00 |
| IgG1_Bisecting | HepatB | 0.492759131 | 7.34E-02 | 1.00E+00 |
| IgG2_Agalactosylation | HepatB | -0.61673081 | 1.88E-02 | 1.00E+00 |
| IgG2_Digalactosylation | HepatB | 0.4293719 | 1.25E-01 | 1.00E+00 |
| IgG2_Sialylation | HepatB | 0.122767741 | 6.76E-01 | 1.00E+00 |
| IgG2_Bisecting | HepatB | 0.368579372 | 1.95E-01 | 1.00E+00 |

| | | | | |
|---|---|---|---|---|
| IgG4_Agalactosylation | HepatB | -0.304482452 | 2.90E-01 | 1.00E+00 |
| IgG4_Monogalactosylation | HepatB | 0.699305953 | 5.38E-03 | 1.00E+00 |
| IgG4_Digalactosylation | HepatB | 0.104314697 | 7.23E-01 | 1.00E+00 |
| IgG4_Sialylation | HepatB | -0.065251782 | 8.25E-01 | 1.00E+00 |
| IgG4_Bisecting | HepatB | 0.156866034 | 5.92E-01 | 1.00E+00 |
| IgG1_Agalactosylation | NTDs | 0.003983452 | 9.89E-01 | 1.00E+00 |
| IgG1_Monogalactosylation | NTDs | 0.025072478 | 9.32E-01 | 1.00E+00 |
| IgG1_Digalactosylation | NTDs | -0.020147237 | 9.45E-01 | 1.00E+00 |
| IgG1_Sialylation | NTDs | -0.046904279 | 8.73E-01 | 1.00E+00 |
| IgG1_Bisecting | NTDs | 0.332299083 | 2.46E-01 | 1.00E+00 |
| IgG2_Agalactosylation | NTDs | 0.095570966 | 7.45E-01 | 1.00E+00 |
| IgG2_Monogalactosylation | NTDs | -0.166334821 | 5.70E-01 | 1.00E+00 |
| IgG2_Digalactosylation | NTDs | -0.10243172 | 7.28E-01 | 1.00E+00 |
| IgG2_Sialylation | NTDs | 0.180646576 | 5.37E-01 | 1.00E+00 |
| IgG2_Bisecting | NTDs | 0.344380425 | 2.28E-01 | 1.00E+00 |
| IgG4_Agalactosylation | NTDs | -0.028134675 | 9.24E-01 | 1.00E+00 |
| IgG4_Monogalactosylation | NTDs | -0.017998302 | 9.51E-01 | 1.00E+00 |
| IgG4_Digalactosylation | NTDs | -0.006793718 | 9.82E-01 | 1.00E+00 |
| IgG4_Sialylation | NTDs | 0.122593774 | 6.76E-01 | 1.00E+00 |
| IgG4_Bisecting | NTDs | 0.205752388 | 4.80E-01 | 1.00E+00 |
| IgG1_Agalactosylation | NCDs | -0.763208648 | 1.49E-03 | 1.00E+00 |
| IgG1_Digalactosylation | NCDs | 0.709659459 | 4.47E-03 | 1.00E+00 |
| IgG1_Sialylation | NCDs | -0.494615449 | 7.22E-02 | 1.00E+00 |
| IgG1_Bisecting | NCDs | 0.611930921 | 2.00E-02 | 1.00E+00 |
| IgG2_Agalactosylation | NCDs | -0.718087966 | 3.82E-03 | 1.00E+00 |
| IgG2_Digalactosylation | NCDs | 0.572247718 | 3.25E-02 | 1.00E+00 |
| IgG2_Sialylation | NCDs | 0.278644748 | 3.35E-01 | 1.00E+00 |
| IgG2_Bisecting | NCDs | 0.533132994 | 4.96E-02 | 1.00E+00 |
| IgG4_Agalactosylation | NCDs | -0.405769059 | 1.50E-01 | 1.00E+00 |
| IgG4_Monogalactosylation | NCDs | 0.679253371 | 7.55E-03 | 1.00E+00 |
| IgG4_Digalactosylation | NCDs | 0.282577889 | 3.28E-01 | 1.00E+00 |
| IgG4_Sialylation | NCDs | -0.01516047 | 9.59E-01 | 1.00E+00 |
| IgG4_Bisecting | NCDs | 0.144924452 | 6.21E-01 | 1.00E+00 |
| IgG1_Agalactosylation | Suicide | -0.28838536 | 3.17E-01 | 1.00E+00 |
| IgG1_Monogalactosylation | Suicide | 0.383192902 | 1.76E-01 | 1.00E+00 |
| IgG1_Digalactosylation | Suicide | 0.281059652 | 3.30E-01 | 1.00E+00 |
| IgG1_Sialylation | Suicide | -0.426197376 | 1.29E-01 | 1.00E+00 |
| IgG1_Bisecting | Suicide | 0.354301464 | 2.14E-01 | 1.00E+00 |
| IgG2_Agalactosylation | Suicide | -0.293529669 | 3.08E-01 | 1.00E+00 |
| IgG2_Monogalactosylation | Suicide | 0.366141401 | 1.98E-01 | 1.00E+00 |
| IgG2_Digalactosylation | Suicide | 0.230680994 | 4.28E-01 | 1.00E+00 |
| IgG2_Sialylation | Suicide | -0.088711598 | 7.63E-01 | 1.00E+00 |
| IgG2_Bisecting | Suicide | 0.397810584 | 1.59E-01 | 1.00E+00 |
| IgG4_Agalactosylation | Suicide | 0.035279648 | 9.05E-01 | 1.00E+00 |
| IgG4_Monogalactosylation | Suicide | 0.191424793 | 5.12E-01 | 1.00E+00 |
| IgG4_Digalactosylation | Suicide | -0.099371818 | 7.35E-01 | 1.00E+00 |
| IgG4_Sialylation | Suicide | -0.247002195 | 3.95E-01 | 1.00E+00 |
| IgG4_Bisecting | Suicide | 0.319975306 | 2.65E-01 | 1.00E+00 |
| IgG1_Agalactosylation | Alcohol | 0.397379917 | 1.59E-01 | 1.00E+00 |
| IgG1_Monogalactosylation | Alcohol | -0.329454176 | 2.50E-01 | 1.00E+00 |
| IgG1_Digalactosylation | Alcohol | -0.317509086 | 2.69E-01 | 1.00E+00 |
| IgG1_Sialylation | Alcohol | -0.083148757 | 7.77E-01 | 1.00E+00 |
| IgG1_Bisecting | Alcohol | -0.257377094 | 3.74E-01 | 1.00E+00 |
| IgG2_Agalactosylation | Alcohol | 0.25818445 | 3.73E-01 | 1.00E+00 |
| IgG2_Monogalactosylation | Alcohol | -0.242742822 | 4.03E-01 | 1.00E+00 |
| IgG2_Digalactosylation | Alcohol | -0.18911913 | 5.17E-01 | 1.00E+00 |
| IgG2_Sialylation | Alcohol | -0.273261169 | 3.45E-01 | 1.00E+00 |
| IgG2_Bisecting | Alcohol | -0.15540424 | 5.96E-01 | 1.00E+00 |
| IgG4_Agalactosylation | Alcohol | 0.382954552 | 1.77E-01 | 1.00E+00 |
| IgG4_Monogalactosylation | Alcohol | -0.374776358 | 1.87E-01 | 1.00E+00 |
| IgG4_Digalactosylation | Alcohol | -0.2912194 | 3.12E-01 | 1.00E+00 |
| IgG4_Sialylation | Alcohol | -0.292786116 | 3.10E-01 | 1.00E+00 |
| IgG4_Bisecting | Alcohol | -0.03195087 | 9.14E-01 | 1.00E+00 |
| IgG1_Agalactosylation | RoadInjuries | -0.724146439 | 3.40E-03 | 1.00E+00 |
| IgG1_Digalactosylation | RoadInjuries | 0.569398699 | 3.36E-02 | 1.00E+00 |
| IgG1_Sialylation | RoadInjuries | -0.310861245 | 2.79E-01 | 1.00E+00 |
| IgG1_Bisecting | RoadInjuries | 0.559478746 | 3.75E-02 | 1.00E+00 |
| IgG2_Agalactosylation | RoadInjuries | -0.467323306 | 9.20E-02 | 1.00E+00 |
| IgG2_Monogalactosylation | RoadInjuries | 0.647845416 | 1.22E-02 | 1.00E+00 |
| IgG2_Digalactosylation | RoadInjuries | 0.27558139 | 3.40E-01 | 1.00E+00 |
| IgG2_Sialylation | RoadInjuries | 0.062064223 | 8.33E-01 | 1.00E+00 |
| IgG2_Bisecting | RoadInjuries | 0.484967013 | 7.88E-02 | 1.00E+00 |
| IgG4_Agalactosylation | RoadInjuries | -0.114322383 | 6.97E-01 | 1.00E+00 |

| | | | | |
|---|---|---|---|---|
| IgG4_Monogalactosylation | RoadInjuries | 0.538160679 | 4.71E-02 | 1.00E+00 |
| IgG4_Digalactosylation | RoadInjuries | -0.09187628 | 7.55E-01 | 1.00E+00 |
| IgG4_Sialylation | RoadInjuries | -0.150732945 | 6.07E-01 | 1.00E+00 |
| IgG4_Bisecting | RoadInjuries | 0.295185425 | 3.06E-01 | 1.00E+00 |
| IgG1_Agalactosylation | FPneedMet | -0.631416341 | 1.54E-02 | 1.00E+00 |
| IgG1_Monogalactosylation | FPneedMet | 0.677482759 | 7.77E-03 | 1.00E+00 |
| IgG1_Digalactosylation | FPneedMet | 0.657751769 | 1.06E-02 | 1.00E+00 |
| IgG1_Sialylation | FPneedMet | -0.485834436 | 7.82E-02 | 1.00E+00 |
| IgG1_Bisecting | FPneedMet | 0.086512634 | 7.69E-01 | 1.00E+00 |
| IgG2_Agalactosylation | FPneedMet | -0.61524581 | 1.92E-02 | 1.00E+00 |
| IgG2_Monogalactosylation | FPneedMet | 0.705505521 | 4.82E-03 | 1.00E+00 |
| IgG2_Digalactosylation | FPneedMet | 0.589614418 | 2.65E-02 | 1.00E+00 |
| IgG2_Sialylation | FPneedMet | -0.054212841 | 8.54E-01 | 1.00E+00 |
| IgG2_Bisecting | FPneedMet | 0.017824051 | 9.52E-01 | 1.00E+00 |
| IgG4_Agalactosylation | FPneedMet | -0.214637897 | 4.61E-01 | 1.00E+00 |
| IgG4_Monogalactosylation | FPneedMet | 0.576068283 | 3.11E-02 | 1.00E+00 |
| IgG4_Digalactosylation | FPneedMet | 0.157424438 | 5.91E-01 | 1.00E+00 |
| IgG4_Sialylation | FPneedMet | -0.275092581 | 3.41E-01 | 1.00E+00 |
| IgG4_Bisecting | FPneedMet | -0.287801317 | 3.18E-01 | 1.00E+00 |
| IgG1_Agalactosylation | AdolBirthRate | -0.740408874 | 2.46E-03 | 1.00E+00 |
| IgG1_Digalactosylation | AdolBirthRate | 0.747298947 | 2.13E-03 | 1.00E+00 |
| IgG1_Sialylation | AdolBirthRate | -0.668919108 | 8.90E-03 | 1.00E+00 |
| IgG1_Bisecting | AdolBirthRate | 0.595527048 | 2.46E-02 | 1.00E+00 |
| IgG2_Agalactosylation | AdolBirthRate | -0.599888648 | 2.33E-02 | 1.00E+00 |
| IgG2_Digalactosylation | AdolBirthRate | 0.559378586 | 3.75E-02 | 1.00E+00 |
| IgG2_Sialylation | AdolBirthRate | -0.183342184 | 5.30E-01 | 1.00E+00 |
| IgG2_Bisecting | AdolBirthRate | 0.533989531 | 4.92E-02 | 1.00E+00 |
| IgG4_Agalactosylation | AdolBirthRate | -0.164733276 | 5.74E-01 | 1.00E+00 |
| IgG4_Monogalactosylation | AdolBirthRate | 0.633020849 | 1.51E-02 | 1.00E+00 |
| IgG4_Digalactosylation | AdolBirthRate | 0.035947295 | 9.03E-01 | 1.00E+00 |
| IgG4_Sialylation | AdolBirthRate | -0.264762893 | 3.60E-01 | 1.00E+00 |
| IgG4_Bisecting | AdolBirthRate | 0.161430215 | 5.81E-01 | 1.00E+00 |
| IgG1_Sialylation | UHCTracer | -0.347857289 | 2.23E-01 | 1.00E+00 |
| IgG1_Bisecting | UHCTracer | 0.596825453 | 2.42E-02 | 1.00E+00 |
| IgG2_Digalactosylation | UHCTracer | 0.611301378 | 2.02E-02 | 1.00E+00 |
| IgG2_Sialylation | UHCTracer | 0.284858144 | 3.24E-01 | 1.00E+00 |
| IgG2_Bisecting | UHCTracer | 0.416957 | 1.38E-01 | 1.00E+00 |
| IgG4_Agalactosylation | UHCTracer | -0.567544362 | 3.43E-02 | 1.00E+00 |
| IgG4_Digalactosylation | UHCTracer | 0.390450897 | 1.68E-01 | 1.00E+00 |
| IgG4_Sialylation | UHCTracer | 0.133111681 | 6.50E-01 | 1.00E+00 |
| IgG4_Bisecting | UHCTracer | 0.043153528 | 8.84E-01 | 1.00E+00 |
| IgG1_Sialylation | AirPollMort | -0.381918984 | 1.78E-01 | 1.00E+00 |
| IgG1_Bisecting | AirPollMort | 0.564891819 | 3.53E-02 | 1.00E+00 |
| IgG2_Agalactosylation | AirPollMort | -0.738456494 | 2.56E-03 | 1.00E+00 |
| IgG2_Digalactosylation | AirPollMort | 0.584565427 | 2.81E-02 | 1.00E+00 |
| IgG2_Sialylation | AirPollMort | 0.163921308 | 5.76E-01 | 1.00E+00 |
| IgG2_Bisecting | AirPollMort | 0.431257539 | 1.24E-01 | 1.00E+00 |
| IgG4_Agalactosylation | AirPollMort | -0.425785871 | 1.29E-01 | 1.00E+00 |
| IgG4_Monogalactosylation | AirPollMort | 0.748217385 | 2.08E-03 | 1.00E+00 |
| IgG4_Digalactosylation | AirPollMort | 0.263719385 | 3.62E-01 | 1.00E+00 |
| IgG4_Sialylation | AirPollMort | 0.029099193 | 9.21E-01 | 1.00E+00 |
| IgG4_Bisecting | AirPollMort | -0.008167816 | 9.78E-01 | 1.00E+00 |
| IgG1_Sialylation | WaSHMort | -0.611132604 | 2.02E-02 | 1.00E+00 |
| IgG2_Agalactosylation | WaSHMort | -0.652927641 | 1.14E-02 | 1.00E+00 |
| IgG2_Monogalactosylation | WaSHMort | 0.746175121 | 2.18E-03 | 1.00E+00 |
| IgG2_Digalactosylation | WaSHMort | 0.596180444 | 2.44E-02 | 1.00E+00 |
| IgG2_Sialylation | WaSHMort | 0.01726152 | 9.53E-01 | 1.00E+00 |
| IgG2_Bisecting | WaSHMort | 0.698411923 | 5.46E-03 | 1.00E+00 |
| IgG4_Agalactosylation | WaSHMort | -0.372968011 | 1.89E-01 | 1.00E+00 |
| IgG4_Monogalactosylation | WaSHMort | 0.687089871 | 6.63E-03 | 1.00E+00 |
| IgG4_Digalactosylation | WaSHMort | 0.235928752 | 4.17E-01 | 1.00E+00 |
| IgG4_Sialylation | WaSHMort | -0.010438687 | 9.72E-01 | 1.00E+00 |
| IgG4_Bisecting | WaSHMort | 0.242987966 | 4.03E-01 | 1.00E+00 |
| IgG1_Agalactosylation | Poisons | -0.648514893 | 1.21E-02 | 1.00E+00 |
| IgG1_Monogalactosylation | Poisons | 0.743496698 | 2.30E-03 | 1.00E+00 |
| IgG1_Digalactosylation | Poisons | 0.597785187 | 2.40E-02 | 1.00E+00 |
| IgG1_Sialylation | Poisons | -0.440526145 | 1.15E-01 | 1.00E+00 |
| IgG1_Bisecting | Poisons | 0.678458832 | 7.64E-03 | 1.00E+00 |
| IgG2_Agalactosylation | Poisons | -0.558160794 | 3.80E-02 | 1.00E+00 |
| IgG2_Monogalactosylation | Poisons | 0.670467914 | 8.69E-03 | 1.00E+00 |
| IgG2_Digalactosylation | Poisons | 0.436428493 | 1.19E-01 | 1.00E+00 |
| IgG2_Sialylation | Poisons | 0.092499433 | 7.53E-01 | 1.00E+00 |
| IgG2_Bisecting | Poisons | 0.582413369 | 2.89E-02 | 1.00E+00 |

| | | | | |
|---|---|---|---|---|
| IgG4_Agalactosylation | Poisons | -0.243672541 | 4.01E-01 | 1.00E+00 |
| IgG4_Monogalactosylation | Poisons | 0.615992284 | 1.90E-02 | 1.00E+00 |
| IgG4_Digalactosylation | Poisons | 0.089943449 | 7.60E-01 | 1.00E+00 |
| IgG4_Sialylation | Poisons | -0.108035979 | 7.13E-01 | 1.00E+00 |
| IgG4_Bisecting | Poisons | 0.189252866 | 5.17E-01 | 1.00E+00 |
| IgG1_Agalactosylation | Smoking | -0.099933896 | 7.34E-01 | 1.00E+00 |
| IgG1_Monogalactosylation | Smoking | -0.046616626 | 8.74E-01 | 1.00E+00 |
| IgG1_Digalactosylation | Smoking | 0.013546408 | 9.63E-01 | 1.00E+00 |
| IgG1_Sialylation | Smoking | 0.362673835 | 2.03E-01 | 1.00E+00 |
| IgG1_Bisecting | Smoking | -0.32246721 | 2.61E-01 | 1.00E+00 |
| IgG2_Agalactosylation | Smoking | -0.166154566 | 5.70E-01 | 1.00E+00 |
| IgG2_Monogalactosylation | Smoking | 0.058853238 | 8.42E-01 | 1.00E+00 |
| IgG2_Digalactosylation | Smoking | 0.041980921 | 8.87E-01 | 1.00E+00 |
| IgG2_Sialylation | Smoking | 0.501186823 | 6.79E-02 | 1.00E+00 |
| IgG2_Bisecting | Smoking | -0.46205343 | 9.62E-02 | 1.00E+00 |
| IgG4_Agalactosylation | Smoking | -0.431116956 | 1.24E-01 | 1.00E+00 |
| IgG4_Monogalactosylation | Smoking | 0.186566743 | 5.23E-01 | 1.00E+00 |
| IgG4_Digalactosylation | Smoking | 0.447683403 | 1.08E-01 | 1.00E+00 |
| IgG4_Sialylation | Smoking | 0.391761623 | 1.66E-01 | 1.00E+00 |
| IgG4_Bisecting | Smoking | -0.399004099 | 1.58E-01 | 1.00E+00 |
| IgG1_Agalactosylation | IPV | -0.419612264 | 1.35E-01 | 1.00E+00 |
| IgG1_Monogalactosylation | IPV | 0.489348659 | 7.57E-02 | 1.00E+00 |
| IgG1_Digalactosylation | IPV | 0.466933299 | 9.23E-02 | 1.00E+00 |
| IgG1_Sialylation | IPV | -0.380704071 | 1.79E-01 | 1.00E+00 |
| IgG1_Bisecting | IPV | 0.21145097 | 4.68E-01 | 1.00E+00 |
| IgG2_Agalactosylation | IPV | -0.233873776 | 4.21E-01 | 1.00E+00 |
| IgG2_Monogalactosylation | IPV | 0.368235122 | 1.95E-01 | 1.00E+00 |
| IgG2_Digalactosylation | IPV | 0.293465971 | 3.09E-01 | 1.00E+00 |
| IgG2_Sialylation | IPV | -0.326295967 | 2.55E-01 | 1.00E+00 |
| IgG2_Bisecting | IPV | 0.160831397 | 5.83E-01 | 1.00E+00 |
| IgG4_Agalactosylation | IPV | 0.078884319 | 7.89E-01 | 1.00E+00 |
| IgG4_Monogalactosylation | IPV | 0.249122518 | 3.90E-01 | 1.00E+00 |
| IgG4_Digalactosylation | IPV | -0.11848905 | 6.87E-01 | 1.00E+00 |
| IgG4_Sialylation | IPV | -0.260186921 | 3.69E-01 | 1.00E+00 |
| IgG4_Bisecting | IPV | -0.090335553 | 7.59E-01 | 1.00E+00 |
| IgG1_Agalactosylation | Water | -0.454339058 | 1.03E-01 | 1.00E+00 |
| IgG1_Monogalactosylation | Water | 0.431752919 | 1.23E-01 | 1.00E+00 |
| IgG1_Digalactosylation | Water | 0.414740925 | 1.40E-01 | 1.00E+00 |
| IgG1_Sialylation | Water | -0.13747466 | 6.39E-01 | 1.00E+00 |
| IgG1_Bisecting | Water | 0.078682325 | 7.89E-01 | 1.00E+00 |
| IgG2_Agalactosylation | Water | -0.33100302 | 2.48E-01 | 1.00E+00 |
| IgG2_Monogalactosylation | Water | 0.318844109 | 2.67E-01 | 1.00E+00 |
| IgG2_Digalactosylation | Water | 0.291060825 | 3.13E-01 | 1.00E+00 |
| IgG2_Sialylation | Water | 0.147575429 | 6.15E-01 | 1.00E+00 |
| IgG2_Bisecting | Water | 0.067696398 | 8.18E-01 | 1.00E+00 |
| IgG4_Agalactosylation | Water | -0.079974915 | 7.86E-01 | 1.00E+00 |
| IgG4_Monogalactosylation | Water | 0.235837663 | 4.17E-01 | 1.00E+00 |
| IgG4_Digalactosylation | Water | 0.034664051 | 9.06E-01 | 1.00E+00 |
| IgG4_Sialylation | Water | -0.108199571 | 7.13E-01 | 1.00E+00 |
| IgG4_Bisecting | Water | -0.059984675 | 8.39E-01 | 1.00E+00 |
| IgG1_Agalactosylation | Sanitation | -0.500133398 | 6.86E-02 | 1.00E+00 |
| IgG1_Monogalactosylation | Sanitation | 0.477524502 | 8.42E-02 | 1.00E+00 |
| IgG1_Digalactosylation | Sanitation | 0.374921373 | 1.87E-01 | 1.00E+00 |
| IgG1_Sialylation | Sanitation | 0.028572446 | 9.23E-01 | 1.00E+00 |
| IgG1_Bisecting | Sanitation | 0.176640513 | 5.46E-01 | 1.00E+00 |
| IgG2_Agalactosylation | Sanitation | -0.352436785 | 2.16E-01 | 1.00E+00 |
| IgG2_Monogalactosylation | Sanitation | 0.36153456 | 2.04E-01 | 1.00E+00 |
| IgG2_Digalactosylation | Sanitation | 0.227472913 | 4.34E-01 | 1.00E+00 |
| IgG2_Sialylation | Sanitation | 0.274291358 | 3.43E-01 | 1.00E+00 |
| IgG2_Bisecting | Sanitation | 0.179309324 | 5.40E-01 | 1.00E+00 |
| IgG4_Agalactosylation | Sanitation | -0.106223199 | 7.18E-01 | 1.00E+00 |
| IgG4_Monogalactosylation | Sanitation | 0.230733157 | 4.27E-01 | 1.00E+00 |
| IgG4_Digalactosylation | Sanitation | 0.005183696 | 9.86E-01 | 1.00E+00 |
| IgG4_Sialylation | Sanitation | 0.001487366 | 9.96E-01 | 1.00E+00 |
| IgG4_Bisecting | Sanitation | 0.027757414 | 9.25E-01 | 1.00E+00 |
| IgG1_Digalactosylation | Hygiene | 0.747485957 | 2.12E-03 | 1.00E+00 |
| IgG1_Sialylation | Hygiene | -0.450870212 | 1.06E-01 | 1.00E+00 |
| IgG1_Bisecting | Hygiene | 0.349683998 | 2.20E-01 | 1.00E+00 |
| IgG2_Agalactosylation | Hygiene | -0.68099141 | 7.34E-03 | 1.00E+00 |
| IgG2_Digalactosylation | Hygiene | 0.579158477 | 3.00E-02 | 1.00E+00 |
| IgG2_Sialylation | Hygiene | 0.031370797 | 9.15E-01 | 1.00E+00 |
| IgG2_Bisecting | Hygiene | 0.282419789 | 3.28E-01 | 1.00E+00 |
| IgG4_Agalactosylation | Hygiene | -0.256146624 | 3.77E-01 | 1.00E+00 |

| | | | | |
|---|---|---|---|---|
| IgG4_Monogalactosylation | Hygiene | 0.666548835 | 9.23E-03 | 1.00E+00 |
| IgG4_Digalactosylation | Hygiene | 0.119961694 | 6.83E-01 | 1.00E+00 |
| IgG4_Sialylation | Hygiene | -0.188565215 | 5.19E-01 | 1.00E+00 |
| IgG4_Bisecting | Hygiene | -0.06214232 | 8.33E-01 | 1.00E+00 |
| IgG1_Agalactosylation | HHAirPoll | -0.720442239 | 3.66E-03 | 1.00E+00 |
| IgG1_Digalactosylation | HHAirPoll | 0.760921273 | 1.57E-03 | 1.00E+00 |
| IgG1_Sialylation | HHAirPoll | -0.532595223 | 4.99E-02 | 1.00E+00 |
| IgG1_Bisecting | HHAirPoll | 0.525572249 | 5.36E-02 | 1.00E+00 |
| IgG2_Agalactosylation | HHAirPoll | -0.63117229 | 1.55E-02 | 1.00E+00 |
| IgG2_Monogalactosylation | HHAirPoll | 0.748424795 | 2.08E-03 | 1.00E+00 |
| IgG2_Digalactosylation | HHAirPoll | 0.631430596 | 1.54E-02 | 1.00E+00 |
| IgG2_Sialylation | HHAirPoll | -0.13154409 | 6.54E-01 | 1.00E+00 |
| IgG2_Bisecting | HHAirPoll | 0.451423307 | 1.05E-01 | 1.00E+00 |
| IgG4_Agalactosylation | HHAirPoll | -0.25040777 | 3.88E-01 | 1.00E+00 |
| IgG4_Monogalactosylation | HHAirPoll | 0.582940986 | 2.87E-02 | 1.00E+00 |
| IgG4_Digalactosylation | HHAirPoll | 0.15971854 | 5.85E-01 | 1.00E+00 |
| IgG4_Sialylation | HHAirPoll | -0.114732718 | 6.96E-01 | 1.00E+00 |
| IgG4_Bisecting | HHAirPoll | -0.10805427 | 7.13E-01 | 1.00E+00 |
| IgG1_Sialylation | OccRiskBurden | -0.310366031 | 2.80E-01 | 1.00E+00 |
| IgG1_Bisecting | OccRiskBurden | 0.636037041 | 1.45E-02 | 1.00E+00 |
| IgG2_Digalactosylation | OccRiskBurden | 0.660271401 | 1.02E-02 | 1.00E+00 |
| IgG2_Sialylation | OccRiskBurden | 0.332632221 | 2.45E-01 | 1.00E+00 |
| IgG2_Bisecting | OccRiskBurden | 0.452935795 | 1.04E-01 | 1.00E+00 |
| IgG4_Agalactosylation | OccRiskBurden | -0.634120283 | 1.49E-02 | 1.00E+00 |
| IgG4_Digalactosylation | OccRiskBurden | 0.473461278 | 8.73E-02 | 1.00E+00 |
| IgG4_Sialylation | OccRiskBurden | 0.233319001 | 4.22E-01 | 1.00E+00 |
| IgG4_Bisecting | OccRiskBurden | -0.060478231 | 8.37E-01 | 1.00E+00 |
| IgG1_Agalactosylation | MeanPM25 | -0.352606175 | 2.16E-01 | 1.00E+00 |
| IgG1_Monogalactosylation | MeanPM25 | 0.441458504 | 1.14E-01 | 1.00E+00 |
| IgG1_Digalactosylation | MeanPM25 | 0.22900934 | 4.31E-01 | 1.00E+00 |
| IgG1_Sialylation | MeanPM25 | -0.012648481 | 9.66E-01 | 1.00E+00 |
| IgG1_Bisecting | MeanPM25 | -0.010973801 | 9.70E-01 | 1.00E+00 |
| IgG2_Agalactosylation | MeanPM25 | -0.151209958 | 6.06E-01 | 1.00E+00 |
| IgG2_Monogalactosylation | MeanPM25 | 0.386065459 | 1.73E-01 | 1.00E+00 |
| IgG2_Digalactosylation | MeanPM25 | 0.047932004 | 8.71E-01 | 1.00E+00 |
| IgG2_Sialylation | MeanPM25 | -0.227502754 | 4.34E-01 | 1.00E+00 |
| IgG2_Bisecting | MeanPM25 | -0.031764367 | 9.14E-01 | 1.00E+00 |
| IgG4_Agalactosylation | MeanPM25 | 0.154811764 | 5.97E-01 | 1.00E+00 |
| IgG4_Monogalactosylation | MeanPM25 | 0.175404906 | 5.49E-01 | 1.00E+00 |
| IgG4_Digalactosylation | MeanPM25 | -0.277974539 | 3.36E-01 | 1.00E+00 |
| IgG4_Sialylation | MeanPM25 | -0.21534634 | 4.60E-01 | 1.00E+00 |
| IgG4_Bisecting | MeanPM25 | -0.100472943 | 7.33E-01 | 1.00E+00 |
| IgG1_Agalactosylation | Violece | -0.500811625 | 6.81E-02 | 1.00E+00 |
| IgG1_Monogalactosylation | Violece | 0.632302075 | 1.53E-02 | 1.00E+00 |
| IgG1_Digalactosylation | Violece | 0.427965071 | 1.27E-01 | 1.00E+00 |
| IgG1_Sialylation | Violece | -0.467387133 | 9.20E-02 | 1.00E+00 |
| IgG1_Bisecting | Violece | 0.46685683 | 9.24E-02 | 1.00E+00 |
| IgG2_Agalactosylation | Violece | -0.35769412 | 2.09E-01 | 1.00E+00 |
| IgG2_Monogalactosylation | Violece | 0.483475635 | 7.99E-02 | 1.00E+00 |
| IgG2_Digalactosylation | Violece | 0.239811099 | 4.09E-01 | 1.00E+00 |
| IgG2_Sialylation | Violece | -0.0075278 | 9.80E-01 | 1.00E+00 |
| IgG2_Bisecting | Violece | 0.480446822 | 8.21E-02 | 1.00E+00 |
| IgG4_Agalactosylation | Violece | 0.05151224 | 8.61E-01 | 1.00E+00 |
| IgG4_Monogalactosylation | Violece | 0.359806799 | 2.06E-01 | 1.00E+00 |
| IgG4_Digalactosylation | Violece | -0.179675001 | 5.39E-01 | 1.00E+00 |
| IgG4_Sialylation | Violece | -0.335928661 | 2.40E-01 | 1.00E+00 |
| IgG4_Bisecting | Violece | 0.409614134 | 1.46E-01 | 1.00E+00 |
| IgG1_Agalactosylation | War | -0.059882122 | 8.39E-01 | 1.00E+00 |
| IgG1_Monogalactosylation | War | 0.065298588 | 8.24E-01 | 1.00E+00 |
| IgG1_Digalactosylation | War | 0.029698473 | 9.20E-01 | 1.00E+00 |
| IgG1_Sialylation | War | 0.050331837 | 8.64E-01 | 1.00E+00 |
| IgG1_Bisecting | War | -0.401681394 | 1.55E-01 | 1.00E+00 |
| IgG2_Agalactosylation | War | -0.055065677 | 8.52E-01 | 1.00E+00 |
| IgG2_Monogalactosylation | War | 0.098731137 | 7.37E-01 | 1.00E+00 |
| IgG2_Digalactosylation | War | 0.024849879 | 9.33E-01 | 1.00E+00 |
| IgG2_Sialylation | War | 0.022279441 | 9.40E-01 | 1.00E+00 |
| IgG2_Bisecting | War | -0.443485664 | 1.12E-01 | 1.00E+00 |
| IgG4_Agalactosylation | War | 0.059622545 | 8.40E-01 | 1.00E+00 |
| IgG4_Monogalactosylation | War | 0.07788774 | 7.91E-01 | 1.00E+00 |
| IgG4_Digalactosylation | War | -0.041798468 | 8.87E-01 | 1.00E+00 |
| IgG4_Sialylation | War | -0.194037949 | 5.06E-01 | 1.00E+00 |
| IgG4_Bisecting | War | -0.266306289 | 3.57E-01 | 1.00E+00 |
| IgG1_Sialylation | MDG | -0.559063328 | 3.77E-02 | 1.00E+00 |

| | | | | |
|---|---|---|---|---|
| IgG1_Bisecting | MDG | 0.649044094 | 1.20E-02 | 1.00E+00 |
| IgG2_Agalactosylation | MDG | -0.752689382 | 1.89E-03 | 1.00E+00 |
| IgG2_Digalactosylation | MDG | 0.653583565 | 1.12E-02 | 1.00E+00 |
| IgG2_Sialylation | MDG | 0.09728731 | 7.41E-01 | 1.00E+00 |
| IgG2_Bisecting | MDG | 0.54147241 | 4.55E-02 | 1.00E+00 |
| IgG4_Agalactosylation | MDG | -0.38532966 | 1.74E-01 | 1.00E+00 |
| IgG4_Digalactosylation | MDG | 0.235367786 | 4.18E-01 | 1.00E+00 |
| IgG4_Sialylation | MDG | -0.074162216 | 8.01E-01 | 1.00E+00 |
| IgG4_Bisecting | MDG | 0.074469515 | 8.00E-01 | 1.00E+00 |
| IgG1_Digalactosylation | Non.MDG | 0.763586031 | 1.48E-03 | 1.00E+00 |
| IgG1_Sialylation | Non.MDG | -0.39564521 | 1.61E-01 | 1.00E+00 |
| IgG1_Bisecting | Non.MDG | 0.444662768 | 1.11E-01 | 1.00E+00 |
| IgG2_Agalactosylation | Non.MDG | -0.669538467 | 8.81E-03 | 1.00E+00 |
| IgG2_Digalactosylation | Non.MDG | 0.541265208 | 4.56E-02 | 1.00E+00 |
| IgG2_Sialylation | Non.MDG | 0.072354456 | 8.06E-01 | 1.00E+00 |
| IgG2_Bisecting | Non.MDG | 0.319796199 | 2.65E-01 | 1.00E+00 |
| IgG4_Agalactosylation | Non.MDG | -0.288542102 | 3.17E-01 | 1.00E+00 |
| IgG4_Monogalactosylation | Non.MDG | 0.684048278 | 6.98E-03 | 1.00E+00 |
| IgG4_Digalactosylation | Non.MDG | 0.139539727 | 6.34E-01 | 1.00E+00 |
| IgG4_Sialylation | Non.MDG | -0.109674342 | 7.09E-01 | 1.00E+00 |
| IgG4_Bisecting | Non.MDG | -0.063397227 | 8.30E-01 | 1.00E+00 |
| IgG1_Sialylation | LifeExp | -0.662663994 | 9.80E-03 | 1.00E+00 |
| IgG1_Bisecting | LifeExp | 0.614494663 | 1.94E-02 | 1.00E+00 |
| IgG2_Agalactosylation | LifeExp | -0.71684363 | 3.91E-03 | 1.00E+00 |
| IgG2_Digalactosylation | LifeExp | 0.655658929 | 1.09E-02 | 1.00E+00 |
| IgG2_Sialylation | LifeExp | 0.020279666 | 9.45E-01 | 1.00E+00 |
| IgG2_Bisecting | LifeExp | 0.560632817 | 3.70E-02 | 1.00E+00 |
| IgG4_Agalactosylation | LifeExp | -0.267980642 | 3.54E-01 | 1.00E+00 |
| IgG4_Monogalactosylation | LifeExp | 0.666783746 | 9.20E-03 | 1.00E+00 |
| IgG4_Digalactosylation | LifeExp | 0.15831053 | 5.89E-01 | 1.00E+00 |
| IgG4_Sialylation | LifeExp | -0.192767766 | 5.09E-01 | 1.00E+00 |
| IgG4_Bisecting | LifeExp | 0.076453001 | 7.95E-01 | 1.00E+00 |
| IgG1_Sialylation | LifeExp_F | -0.662438241 | 9.84E-03 | 1.00E+00 |
| IgG1_Bisecting | LifeExp_F | 0.66832596 | 8.98E-03 | 1.00E+00 |
| IgG2_Agalactosylation | LifeExp_F | -0.749277696 | 2.04E-03 | 1.00E+00 |
| IgG2_Digalactosylation | LifeExp_F | 0.695195443 | 5.78E-03 | 1.00E+00 |
| IgG2_Sialylation | LifeExp_F | 0.039962836 | 8.92E-01 | 1.00E+00 |
| IgG2_Bisecting | LifeExp_F | 0.607084012 | 2.13E-02 | 1.00E+00 |
| IgG4_Agalactosylation | LifeExp_F | -0.325427159 | 2.56E-01 | 1.00E+00 |
| IgG4_Monogalactosylation | LifeExp_F | 0.69788165 | 5.51E-03 | 1.00E+00 |
| IgG4_Digalactosylation | LifeExp_F | 0.214268566 | 4.62E-01 | 1.00E+00 |
| IgG4_Sialylation | LifeExp_F | -0.136057774 | 6.43E-01 | 1.00E+00 |
| IgG4_Bisecting | LifeExp_F | 0.070999032 | 8.09E-01 | 1.00E+00 |
| IgG1_Agalactosylation | LifeExp_M | -0.759156802 | 1.64E-03 | 1.00E+00 |
| IgG1_Digalactosylation | LifeExp_M | 0.75944531 | 1.63E-03 | 1.00E+00 |
| IgG1_Sialylation | LifeExp_M | -0.64849518 | 1.21E-02 | 1.00E+00 |
| IgG1_Bisecting | LifeExp_M | 0.553120978 | 4.02E-02 | 1.00E+00 |
| IgG2_Agalactosylation | LifeExp_M | -0.678035087 | 7.70E-03 | 1.00E+00 |
| IgG2_Digalactosylation | LifeExp_M | 0.608383469 | 2.10E-02 | 1.00E+00 |
| IgG2_Sialylation | LifeExp_M | 0.01312646 | 9.64E-01 | 1.00E+00 |
| IgG2_Bisecting | LifeExp_M | 0.505541209 | 6.52E-02 | 1.00E+00 |
| IgG4_Agalactosylation | LifeExp_M | -0.218670542 | 4.53E-01 | 1.00E+00 |
| IgG4_Monogalactosylation | LifeExp_M | 0.63055583 | 1.56E-02 | 1.00E+00 |
| IgG4_Digalactosylation | LifeExp_M | 0.112452061 | 7.02E-01 | 1.00E+00 |
| IgG4_Sialylation | LifeExp_M | -0.236036038 | 4.17E-01 | 1.00E+00 |
| IgG4_Bisecting | LifeExp_M | 0.083298089 | 7.77E-01 | 1.00E+00 |
| IgG1_Digalactosylation | GDP_2013 | 0.722327529 | 3.53E-03 | 1.00E+00 |
| IgG1_Sialylation | GDP_2013 | -0.370525677 | 1.92E-01 | 1.00E+00 |
| IgG1_Bisecting | GDP_2013 | 0.396319233 | 1.61E-01 | 1.00E+00 |
| IgG2_Agalactosylation | GDP_2013 | -0.660469461 | 1.01E-02 | 1.00E+00 |
| IgG2_Digalactosylation | GDP_2013 | 0.507510019 | 6.40E-02 | 1.00E+00 |
| IgG2_Sialylation | GDP_2013 | 0.041486819 | 8.88E-01 | 1.00E+00 |
| IgG2_Bisecting | GDP_2013 | 0.265866083 | 3.58E-01 | 1.00E+00 |
| IgG4_Agalactosylation | GDP_2013 | -0.294156887 | 3.07E-01 | 1.00E+00 |
| IgG4_Monogalactosylation | GDP_2013 | 0.726472538 | 3.25E-03 | 1.00E+00 |
| IgG4_Digalactosylation | GDP_2013 | 0.121055593 | 6.80E-01 | 1.00E+00 |
| IgG4_Sialylation | GDP_2013 | -0.136002315 | 6.43E-01 | 1.00E+00 |
| IgG4_Bisecting | GDP_2013 | -0.034072452 | 9.08E-01 | 1.00E+00 |
| IgG1_Sialylation | EduIndx_2014 | -0.383263188 | 1.76E-01 | 1.00E+00 |
| IgG1_Bisecting | EduIndx_2014 | 0.659262122 | 1.03E-02 | 1.00E+00 |
| IgG2_Agalactosylation | EduIndx_2014 | -0.695746356 | 5.72E-03 | 1.00E+00 |
| IgG2_Digalactosylation | EduIndx_2014 | 0.553948047 | 3.98E-02 | 1.00E+00 |
| IgG2_Sialylation | EduIndx_2014 | 0.206766445 | 4.78E-01 | 1.00E+00 |

| | | | | |
|---|---|---|---|---|
| IgG2_Bisecting | EduIndx_2014 | 0.527698 | 5.25E-02 | 1.00E+00 |
| IgG4_Agalactosylation | EduIndx_2014 | -0.408763064 | 1.47E-01 | 1.00E+00 |
| IgG4_Digalactosylation | EduIndx_2014 | 0.225728967 | 4.38E-01 | 1.00E+00 |
| IgG4_Sialylation | EduIndx_2014 | 0.016480716 | 9.55E-01 | 1.00E+00 |
| IgG4_Bisecting | EduIndx_2014 | 0.096558068 | 7.43E-01 | 1.00E+00 |
| IgG1_Sialylation | HealthIndx_2014 | -0.662358154 | 9.85E-03 | 1.00E+00 |
| IgG1_Bisecting | HealthIndx_2014 | 0.615823045 | 1.90E-02 | 1.00E+00 |
| IgG2_Agalactosylation | HealthIndx_2014 | -0.716725057 | 3.92E-03 | 1.00E+00 |
| IgG2_Digalactosylation | HealthIndx_2014 | 0.655358606 | 1.10E-02 | 1.00E+00 |
| IgG2_Sialylation | HealthIndx_2014 | 0.020945002 | 9.43E-01 | 1.00E+00 |
| IgG2_Bisecting | HealthIndx_2014 | 0.562134208 | 3.64E-02 | 1.00E+00 |
| IgG4_Agalactosylation | HealthIndx_2014 | -0.268198259 | 3.54E-01 | 1.00E+00 |
| IgG4_Monogalactosylation | HealthIndx_2014 | 0.666583879 | 9.23E-03 | 1.00E+00 |
| IgG4_Digalactosylation | HealthIndx_2014 | 0.158356062 | 5.89E-01 | 1.00E+00 |
| IgG4_Sialylation | HealthIndx_2014 | -0.192002517 | 5.11E-01 | 1.00E+00 |
| IgG4_Bisecting | HealthIndx_2014 | 0.077687209 | 7.92E-01 | 1.00E+00 |
| IgG1_Sialylation | Water_UN | -0.379683953 | 1.81E-01 | 1.00E+00 |
| IgG2_Monogalactosylation | Water_UN | 0.751575387 | 1.94E-03 | 1.00E+00 |
| IgG2_Digalactosylation | Water_UN | 0.741306579 | 2.41E-03 | 1.00E+00 |
| IgG2_Sialylation | Water_UN | 0.556240275 | 3.89E-02 | 1.00E+00 |
| IgG2_Bisecting | Water_UN | 0.59416513 | 2.51E-02 | 1.00E+00 |
| IgG4_Digalactosylation | Water_UN | 0.704510245 | 4.91E-03 | 1.00E+00 |
| IgG4_Sialylation | Water_UN | 0.387514548 | 1.71E-01 | 1.00E+00 |
| IgG4_Bisecting | Water_UN | -0.043416056 | 8.83E-01 | 1.00E+00 |
| IgG1_Sialylation | Sanitation_UN | -0.507815664 | 6.38E-02 | 1.00E+00 |
| IgG1_Bisecting | Sanitation_UN | 0.728398612 | 3.13E-03 | 1.00E+00 |
| IgG2_Digalactosylation | Sanitation_UN | 0.713336362 | 4.18E-03 | 1.00E+00 |
| IgG2_Sialylation | Sanitation_UN | 0.238386729 | 4.12E-01 | 1.00E+00 |
| IgG2_Bisecting | Sanitation_UN | 0.59228991 | 2.56E-02 | 1.00E+00 |
| IgG4_Agalactosylation | Sanitation_UN | -0.520945372 | 5.61E-02 | 1.00E+00 |
| IgG4_Digalactosylation | Sanitation_UN | 0.409075318 | 1.46E-01 | 1.00E+00 |
| IgG4_Sialylation | Sanitation_UN | 0.101216901 | 7.31E-01 | 1.00E+00 |
| IgG4_Bisecting | Sanitation_UN | -0.057322838 | 8.46E-01 | 1.00E+00 |
